# Simulated digestion, uptake and colonic fermentation of Gorse (*Ulex europeaus*) suggests safe use as a source of protein concentrates and bioactives

**DOI:** 10.1101/2023.11.30.569454

**Authors:** Ajay Iyer, Gary J. Duncan, Nicholas J. Hayward, Charles S. Bestwick, Sylvia H. Duncan, Wendy R. Russell

## Abstract

**Background:** Invasive plants such as Gorse (*Ulex europeaus*) may serve as a potentially sustainable source of edible proteins and bioactives through the production of leaf protein concentrates (GLPC). The digestion, uptake and safety of phytochemicals from GLPC and the protein depleted but bioactive rich fraction (PDF) from Gorse was studied using an in vitro simulated digestion model.

**Results:** Both the Gorse fractions and their digesta maintained the viability of cultured (Caco-2) intestinal cells above the 80% threshold, suggesting a lack of overt toxicity.

In the digestion model, most of the plant metabolites in the digesta were largely confined to the apical fraction of the Caco-2 monolayer (representing the lumen) and were subsequently expected to be delivered to the colon. Food matrix had a significant but marginal effect on the permeability of metabolites across the Caco-2 monolayer. Lastly, *in vitro* intestinal microbial action appeared to produce beneficial compounds such as enterolactones, and equol.

**Conclusion:** The Gorse fractions lack general overt toxicity to Caco-2 cells. GLPC and PDF provide metabolic substrates for the colonic microbes to produce secondary compounds that are associated with positive health outcomes.

## 1 INTRODUCTION

The spread of invasive plants can significantly affect local ecologies and subsequently the economy by depreciating the land capability (Vilà & Ibáñez, 2011), competing with crops of agricultural value (Ziska et al., 2011), and disrupting local niches and ecological webs (Cocîrlea & Oancea, 2022). Among the numerous strategies investigated to counter the spread of invasive species, invasivory has been proposed as an unconventional yet sustainable alternative to incineration, animal grazing, or herbicide use (Iyer et al., 2021). Invasivory is the consumption of invasive species as a means to deplete its population in non-native habitats. The consumption of invasive species does not directly compete with the existing agricultural infrastructure and can be easily set up to complement existing food production capacities. In the context of policy-making and urban foraging communities, invasivory has gained interest (Cerveira et al., 2022; Mishan & Hamada, 2020), although formal scientific investigation on this matter remains limited. Gorse has spread far beyond its native habitat that has resulted in substantial ecological and economic damage to the introduced environment (Broadfield & McHenry, 2019). For example, the regions of Canterbury and Wellington in New Zealand are well known cases of Gorse overwhelming local flora and disrupting the ecology (Came et al., 2020).

Previous research by Iyer et al. (2021) has identified Gorse as a lucrative candidate for protein production. During early growth phases, the leaves of Gorse are rich in protein as well as phytochemicals such as chlorogenic acid, kaempferol and sinapic acid. Leaf protein concentrates (LPC) from Gorse offers an unconventional, yet potentially sustainable and scalable way of obtaining nutrition from invasive plant biomass. Gorse leaf protein concentrate (GLPC) can be prepared at a yield and purity of 5.5% (w·w^−1^) and 96.8% (w·w^−1^) respectively, with a highly favourable amino acid profile (Iyer et al., 2022). While LPC has previously been studied in a human intervention setup (Dewan et al., 2007), it was important to establish the safety of GLPC for application in consumer products.

Establishing food safety in the preliminary laboratory stages of product development involves establishing the absence of potent anti-nutrients, and a precedence of consumption, according to the SAFEST guidelines (Jonas et al., 1996). As of 23 August 2023, no reports of toxicity *via* anti-nutrients in Gorse could be found on literature databases such as The National Library of Medicine (MEDLINE), Cochrane Central, Scopus and Google Scholar. Conversely though, there is limited evidence of its direct human consumption, with the bulk of the documentation largely focused on animal feed. Gorse continues to remain a subsistence form of animal feed when high quality feeds are unavailable (Rymer, 1979).

The aim of the study was to test if GLPC and plant metabolites were safe for consumption. In this work, GLPC and the protein depleted fractions (PDF), obtained from Gorse leaves, were subjected to an in vitro simulated digestion model to understand their digestion as they progress through the digestive tract. The uptake of metabolites at the intestinal phase in the model was also simulated using polarised Caco-2 cells. Additionally, the cell viability assays using the Caco-2 cell model (at their 36 hour proliferative stage) was used to test for the absence of toxicity.

## 2 MATERIALS AND METHODS

All chemicals were purchased from Merck (Darmstadt, Germany) unless stated otherwise. Milli-Q^®^purified water was used for all experiments. To keep the media and water anaerobic, they were boiled for 5 min, allowed to cool for one hour, then flushed with nitrogen for 2 min. Finally, the water was placed in an anaerobic cabinet (Don Whitley Scientific, West Yorkshire, UK) with a gas phase of 10% H_2_, 10% CO_2_, and 80% N_2_, for 48 h before use.

### 2.1 GORSE SAMPLE PREPARATION

Gorse (*Ulex europaeus*) was collected in Aberdeenshire, Scotland (GPS co-ordinate: 57.257, −2.483). Sampling procedures and processing of the Gorse leaves has been previously described by Iyer et al. (2022).

The freeze-dried and milled Gorse leaves (20 g) were first macerated in phosphate buffer solution (PBS; 20 mM; pH 7.0; 200 mL) at 30 °C for 30 min. The sample was centrifuged (3000 × *g* ; 15 min) and the supernatant was recovered and set aside. The retentate was subjected to a second extraction using cellulase (1 mg·g^−1^ dry retentate) in sodium-citrate buffer (50 mM; pH 4.0; 200 mL) at 40 °C for 2 h. The sample was centrifuged (3000 × *g* ; 15 min) and the supernatant was pooled with the previous step. Ethanol (130 mL) was added to the pooled supernatants and allowed to rest at −20 °C overnight. The sample was centrifuged (4000 × *g* ; 30 min). The supernatant (after *in vacuo* ethanol removal) and precipitate were freeze-dried. The mass obtained from dried supernatant was the protein depleted fraction (PDF) and the precipitate was the leaf protein concentrate (GLPC).

### 2.2 THE SIMULATED DIGESTION MODEL

The *in vitro* digestion model used to study the GLPC and PDF samples was based on a previous publication by Minekus et al. (2014) with some modifications using other protocols by (Mulet-Cabero et al., 2020; Brodkorb et al., 2019).

Stock solutions (2.5*×*, 200 mL) were prepared as follows:

Stock salivary fluid comprised of KCl (18.87 mM), KH_2_PO_4_ (4.63 mM), NaHCO_3_ (17.0 mM), MgCl_2_ *·* 6 H_2_O (0.18 mM) and (NH_4_)_2_CO_3_ (0.08 mM).

Stock gastric fluid comprised of KCl (8.62 mM), KH_2_PO_4_ (1.13 mM), NaHCO_3_ (31.25 mM), NaCl (9.6 mM), MgCl_2_ *·* 6 H_2_O (0.41 mM) and (NH_4_)_2_CO_3_ (0.63 mM).

Stock intestinal fluid comprised of KCl (8.5 mM), KH_2_PO_4_ (1.0 mM), NaHCO_3_ (106.25 mM), NaCl (48.0 mM), MgCl_2_ *·* 6 H_2_O (0.41 mM) and (NH_4_)_2_CO_3_ (0.41 mM).

Calcium chloride (CaCl_2_ *·* 2 H_2_O) solution (10 mL, 300.0 mM) was also prepared which was added to the final simulated fluids.

Enzyme solutions *α*-amylase (0.15 g·mL^−1^; Merck, Darmstast, Germany), pepsin (0.1 g·mL^−1^; *≥*30 units·mg Merck, Darmstadt, Germany) and pancreatin (0.08 g·mL^−1^; Merck, Darmstadt, Germany) were prepared using the stock salivary, gastric, and intestinal buffers respectively. Thus the final experimental Simulated Salivary Fluid (SSF) comprised of stock salivary fluid (70 mL), *α*-amylase solution (10 mL), calcium chloride solution (CaCl, 0.48 mL) with pH adjusted to 7 using sodium hydroxide solution (NaOH, 1 M). Simulated Gastric Fluid (SGF) comprised of stock gastric fluid (64 mL), pepsin solution (16 mL), calcium chloride solution (CaCl, 0.48 mL) and adjusted to pH 4 using hydrochloric acid solution (HCl, 1 M). Simulated Intestinal Fluid (SIF) comprised of stock intestinal fluid (42.5 mL), pancreatin solution (25 mL), calcium chloride solution (CaCl, 0.2 mL) and adjusted to pH 7 using sodium hydroxide solution (NaOH, 1 M).

All simulated fluids were maintained at 37 °C for at least 20 min prior to sample addition. SGF and SIF were prepared and stored in an anaerobic cabinet. To initiate the simulated salivary phase, GLPC/PDF (2 g) was incubated with SSF (20 mL; 37 °C) for 30 s, while stirring on a magnetic stirrer. The relative volume and incubation period was adjusted according to physiological reports (Hughes & Wiles, 1996). The sample and SSF were then transferred to an anaerobic chamber (37 °C) where they were added to SGF (40 mL) and incubated under stirring for 3 h. Samples were finally transferred to the SIF (80 mL) and incubated for a further 3 h while stirring. These incubation times were based on physiological transit periods in an empty alimentary tract (Schiller et al., 2005). Samples were then centrifuged at 3000 × *g* for 15 min and the supernatant was sterilised using 0.2 µm filter. At each stage, 1 mL of sample was taken for the analysis of phenolics *via* LC/MS.

### 2.3 CELL CULTURE

#### 2.3.1 MEDIA

The complete working medium (CWM) was comprised of DMEM (863.6 mL·L^−1^; Dulbecco’s Modified Eagle Medium; BioWhittaker^®^; Lonza; Walkersville; USA), foetal Bovine Serum (104.9 mL; HyClone^™^; GE Healthcare Life Sciences; Utah; USA), penicillin-streptomycin 100*×* solution (10.5 mL·L^−1^; Sigma-Aldrich, Pennsylvania; USA) and non-essential amino acids (10.5 mL·L^−1^; Sigma-Aldrich, Pennsylvania; USA).

Phenol-red free complete working medium (PRF-CWM) was comprised of DMEM (874.1 mL·L^−1^; DMEM; BioWhittaker^®^; Lonza; Walkersville; USA), foetal bovine serum (104.9 mL·L^−1^; HyClone^™^; GE Healthcare Life Sciences; Utah; USA), penicillin-streptomycin 100*×* solution (10.5 mL·L^−1^; Sigma-Aldrich, Pennsylvania; USA), non-essential amino acids (NEAA, 10.5 mL·L^−1^, Sigma-Aldrich, Pennsylvania, USA) and glutamine (10.5 mL·L^−1^; Sigma-Aldrich, Pennsylvania; USA).

#### 2.3.2 CELL TYPE AND HANDLING

Caco-2 cells (at passage number 21) were purchased from ATCC and maintained at −70 °C under glycerol in liquid nitrogen at a concentration of 10^6^ cells·mL^−1^. Basic handling and set up were performed based on established good laboratory practices Natoli et al., 2012. The working culture was adjusted to 10^4^ cells·mL^−1^ and incubated in 10 mL CWM. Media was replaced every 48 h.

### 2.4 SAMPLE AND STANDARD PREPARATION

#### 2.4.1 REPRESENTATIVE/STANDARD COMPOUNDS

In a previous publication by Iyer et al. (2021), ten compounds were found to be significantly higher in Gorse compared to other invasive plants found in Scotland, namely, p-coumaric acid, acetovanillone, protocatechuic acid, chlorogenic acid, vanillin, p-hydroxybenzoic acid, sinapic acid, kaempferol, mandelic acid and ferulic acid. Among these however, only six compounds, namely, p-coumaric acid, acetovanillone, vanillin, p-hydroxybenzoic acid, mandelic acid and ferulic acid were detected in the final GLPC and were used as representative sample compounds to study their metabolism as they passed along the digestion model. A standard compound mixture (50 mM, 200 µL, p-coumaric acid, acetovanillone, protocatechuic acid, chlorogenic acid, vanillin, p-hydroxybenzoic acid, sinapic acid, kaempferol, mandelic acid and ferulic acid) was prepared and incubated with the Caco-2 cells for 24 h at 37 °C to assess active cell transport.

#### 2.4.2 PREPARATION OF GLPC AND PDF DIGESTA FOR TOXICITY TESTING

Samples obtained at the intestinal phase in the digestion model were collected, freeze-dried and reconstituted in PRF-CWM. The samples were filter-sterilised through a 0.2 µm filter (Millipore, USA). Toxicity was tested on these samples as described above.

### 2.5 TOXICITY

#### 2.5.1 MTT ASSAY

A sterile 96-well plate (Greiner Bio-One, Kremsmünster, Austria) was seeded with Caco-2 cells (6×10^4^ cells; 200 µL), which were allowed to adhere to the plate over 36 h. The spent media was aspirated, and cells were washed *in situ* using sterile PBS. Then the digestate samples (200 µL) were loaded and incubated for 24 h. The supernatant medium was drained, and the cells were incubated in PRF-CWM (100 µL) with MTT (3-(4,5-dimethylthiazol-2-yl)-2,5-diphenyltetrazolium bromide) at a concentration of 0.5 mg·mL^−1^ for 4 h at 37 °C. The supernatant was aspirated and acidified (40 mM, HCl), and 2-propanol (250 µL) was added to the wells. The 96-well plate was placed on a shaker for 30 min, then centrifuged at 900 × *g* at 4 °C for 10 min min. The supernatant (150 µL) was recovered and absorbance was measured at 570 nm.

### 2.6 METABOLITE TRANSPORT ACROSS THE CACO-2 MONOLAYER

The transport of metabolites in the digestates across the intestinal barrier was simulated using a Caco-2 monolayer on a transwell plate (Hubatsch et al., 2007). Metabolite movement was measured in both directions across the Caco-2 monolayer to establish if transportation was passive or active, and independent or synergistic/ inhibitory. The basolateral fraction represented the blood where metabolites were expected to be measured after being transported across the Caco-2 monolayer. The apical fraction represented the lumen, where metabolites which could not be transported across the Caco-2 monolayer were expected to remain. Under physiological conditions, the apical (luminal) fraction was expected to be delivered to the colon where it interacts with commensal microbes.

#### 2.6.1 APICAL TO BASOLATERAL

Two 24-well plates with filter-inserts (0.4 µm pore size; Greiner Bio-One, Kremsmünster, Austria) were seeded with Caco-2 (10^4^ cells·cm^−2^ at passage 27) and incubated for 21 d to facilitate growth and differentiation. The Transepithelial Electrical Resistance (TEER) was measured to be 4879±360 mΩ. Once TEER measurements were found to be constant and consistent across wells, PDF, GLPC, or standard compound (200 µL) was loaded on the upper well. The lower well contained PRF-CWM (1 mL) which was drawn hourly for 4 h and stored immediately at −20 °C. The bottom well was replenished with fresh PRF-CWM with every draw of sample.

#### 2.6.2 BASOLATERAL TO APICAL

In a technique described previously by Stone et al. (2019), wells were initially layered with a poly-L-lysine solution (Sigma-Aldrich, Pennsylvania, USA) for 2 h and allowed to air-dry overnight in sterile conditions. The inserts were inverted, arranged on the lid, and seeded with Caco-2 (10^4^ cells·cm^−2^, passage 38). The baseplate was gently laid over the inserts as the bottom chamber touched the seed droplet, stabilising them as a hanging droplet. The set-up was allowed to rest at 37 °C for 6 h. A fresh 24-well plate was filled with PRF-CWM (1 mL) and the inserts were gently laid into them and incubated for two weeks to facilitate growth and differentiation. TEER was measured to be 5055±666 mΩ. The subsequent steps were identical to the apical to basolateral part of the experiment.

### 2.7 SCALED UP METABOLITE TRANSPORT

The apical to basolateral transport was scaled up using a 100 mm diameter culture dish (Corning, Transwell^®^, 75 mm insert, polycarbonate, 44 cm^2^ surface area) under conditions described above. The TEER was 454±92 mΩ.

### 2.8 FAECAL FERMENTATION

#### 2.8.1 MEDIUM

The M2GSC medium was prepared as described previously by Cummings and Macfarlane (1991). The recipe is as follows (1*×* concentration):

Bacto^™^Casitone (10 g·L^−1^), Bacto^™^Yeast Extract (2.5 g·L^−1^), sodium bicarbonate (NaHCO_3_, 4.0 g·L^−1^), glucose (2 g·L^−1^), soluble starch (2 g·L^−1^), cellobiose (2 g·L^−1^), clarified rumen fluid (100 mL·L^−1^), cysteine HCl (1 g·L^−1^), dipotassium hydrogen phosphate (K_2_HPO_4_, 0.4 g·L^−1^), potassium dihydrogen phosphate (K_2_HPO_4_, 0.4 g·L^−1^), ammonium sulphate ((NH_4_)_2_SO_4_, 0.8 g·L^−1^), sodium chloride (NaCl, 0.8 g·L^−1^), magnesium sulphate (MgSO_4_, 80 mg·L^−1^), calcium chloride (CaCl_2_, 80 mg·L^−1^) and resazurin (1 µg·L^−1^).

The solution was prepared using water kept under anaerobic conditions for 48 h.

#### 2.8.2 FERMENTATION SAMPLE PREPARATION

The luminal fraction of GLPC and PDF digesta from the preceding Transwell^®^setup were recovered and freeze-dried. The samples were reconstituted in M2GSC medium and filter sterilised under anaerobic conditions to produce the experimental medium for faecal fermentation.

#### 2.8.3 FAECAL INOCULATION

Faecal samples were obtained from three healthy adult donors, stored at 4 °C, and processed within 5 h of collection. Under sterile conditions, the faecal sample from each donor (2 g) was weighed into DispoMix^™^tubes (gentleMACS) containing sterile glycerol-PBS (PBS in 30% glycerol, deoxygenated and autoclaved, 10 mL). The samples were blended under high shear using the DispoMix^™^blender (Medic Tools; Switzerland) for 30 s to obtain faecal slurries.

Experimental medium (5 mL), and faecal slurry (20 µL) were suspended in air-tight, rubber-sealed tubes and incubated at 37 °C for 48 h. Samples were drawn using anaerobic techniques at 0 h, 24 h and 48 h and stored at −70 °C for no longer than 20 d until further processing. For further analysis, samples were allowed to thaw overnight at 4 °C in the dark and centrifuged at 12,500 × *g* for 5 min at 4 °C. The supernatant was recovered and characterised for metabolites.

### 2.9 METABOLITE CHARACTERISATION

Phenolic and protein metabolites present in the analytes were characterised using LC-MS as described previously by Russell et al. (2011). Naming schemes used for in house measurement, corresponding trivial names, and the compounds groups are available in **Supplementary Table 1**.

### 2.10 SHORT CHAIN FATTY ACID (SCFA) MEASUREMENTS

Short chain fatty acids were measured using gas chromatography as described by Richardson et al. (1989). Internal standard used was EBA (50 µL; 2-ethyl butyric acid; 1.27% v·v^−1^). External working standard; prepared in deoxygenated distilled water, comprised of acetic acid (1.72 mL·L^−1^, Fisher, UK), propionic acid (1.49 mL·L^−1^), iso-butyric acid (0.5 mL·L^−1^), n-butyric acid (0.18 mL·L^−1^), sodium formate (0.7 g·L^−1^, BDH Chemicals Ltd., UK), lithium lactate (1.0 g·L^−1^, BDH Chemicals Ltd., UK), and sodium succinate dibasic hexahydrate (0.3 g·L^−1^).

External standard or sample (1 mL) was mixed with HCl (hydrochloric acid, 0.5 mL, 9 M) and diethyl ether (2 mL) using a vortex mixer for one minute. The mixture was centrifuged at 3000 × *g* for 10 min under ambient condition. The organic layer was recovered and to the aqueous fraction, fresh diethyl ether (2 mL) was added and subjected to extraction once more. The organic layers were pooled. From the pooled organic layer, 800 µL sample and MTBSTFA (100 µL, N-tert-butyldimethylsilyl)-N-methyltrifluoroacetamide) were transferred to amber capped glass vials (Agilent Technologies, USA), heated to 80 °C for 20 min. The vials were allowed to rest under ambient conditions in the dark for 72 h.

Derivatised SCFA samples were measured on an Agilent HP 6890 GC system with autosampler and cooled sample tray (between 8 to 12 °C), using an Agilent 190912-333 HP-1: Methyl-Siloxane column. Samples were heated to 63 °C for 3 min and held at this temperature for a further 3 min. Along a temperature ramp of 10 °C·min^−1^, column temperature was increased to 190 °C within 17.2 min, and held for 1.5 min. The evaporated sample was further heated to 275 °C and maintained at 135.0 mN·mm^−2^ within the GC. The measurement was performed using Flame Ionization Detector unit (FID).

Owing to constraints in available resources during the Covid-19 pandemic, the triplicate samples generated from the experiment were pooled into a single sample and measured for SCFA.

### 2.11 STATISTICAL ANALYSIS

Statistical analysis was performed using R (Version 3.6.3) (R Core Team, 2021). Data processing and visualisation were performed using the tidyverse set of packages (Wickham et al., 2019). Principal component analysis (PCA, univariate-scaled) was performed using packages factoextra (Kassambara & Mundt, 2020). Statistical comparison across means was performed using one-way ANOVA and expressed as F_(degrees of freedom, residuals)_=F value, p-value). The raw data, codes and annotations may be found in the OSF repository (https://osf.io/9qtyz/).

Three univariate-scaled principal component analyses (PCA) were performed on the data set. The first PCA model considered the non-bacterial phase of the model that included the metabolite profile of the initial GLPC/PDF fractions to the apical and basolateral fractions after Caco-2 exposure. The second PCA model considered the apical fraction of the Caco-2 model to the 48 h simulated colonic fermentation. The third PCA model considered the complete set of phases from the GLPC/PDF to the fermentation phase.

## 3 RESULTS

### 3.1 METABOLITES ACROSS THE *IN VITRO* DIGESTION MODEL

The LC/MS platform detected and quantified 181 metabolites in the *in vitro* digestion model. Data for the metabolites measured in each phase of the digestion model, including control measurements, is provided in **Supplementary Tables 2** and **3**. **Supplementary Table 2** presents the metabolites quantified for samples from the initial GLPC/PDF to the Caco-2 monolayer. **Supplementary Table 3** presents metabolites quantified in the faecal fermentation. Corresponding metabolite group-wise mass balance is provided in **Supplementary Table 4** from the initial GLPC/PDF to the Caco-2 monolayer phase. **Supplementary Table 5** shows the group-wise balance of the faecal fermentation phase of the digestion model.

Interesting dynamics were noted for the major metabolite groups. For mandelic acids in the PDF they could only be detected until the gastric phase (**Supplementary Table 4**), and were subsequently absent even in the supernatants in the faecal fermentation phase (**Supplementary Table 5**). Mandelic acids were absent in the GLPC and its digestates. Similarly in the case of phenolic alcohols, they were absent in GLPC and could only be detected in the PDF until the salivary phase (**Supplementary Table 4**). However, they reappeared at 48 h in the colonic fermentation supernatants (**Supplementary Table 5**). This was expected as many microbes are capable of converting flavonoids into corresponding phenolic alcohols.

For analyses, the *in vitro* digestion model was divided into the non-bacterial (initial GLPC/PDF material to Caco-2 monolayer phase) and bacterial phase (Caco-2 apical fraction to 48 h time point of faecal fermentation), as the metabolites were better represented in the PCA model based on their cos^2^ values.

#### 3.1.1 METABOLITES MEASURED IN GLPC/PDF TO CACO-2 MONOLAYER MODEL

PCA was performed on the data from **Supplementary Table 2**. The codes, and annotations are provided in **Supplementary Code (code chunk 4 and 5)**. This model captures the variance in metabolites from GLPC/PDF to the Caco-2 monolayer, and is depicted in Figure 1.

**Figure 1:**
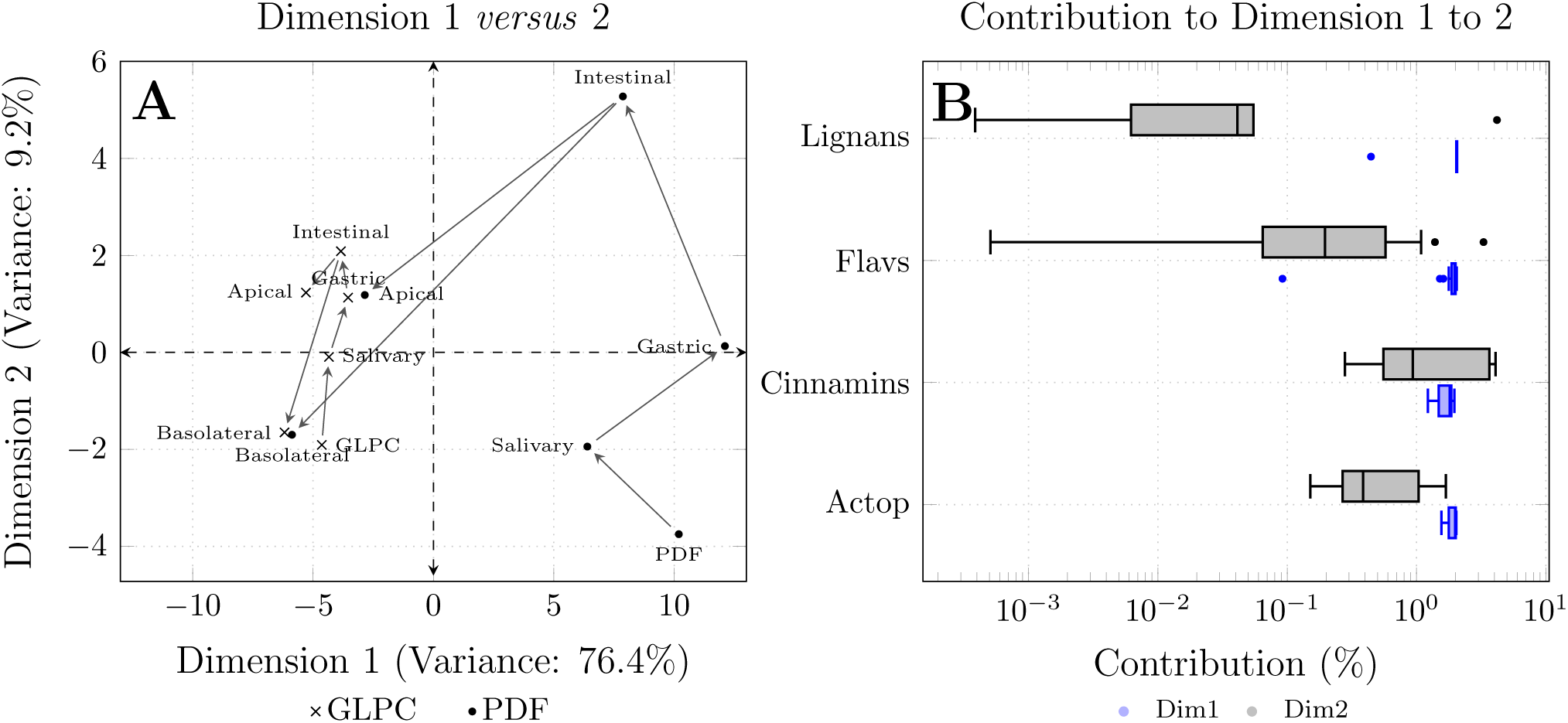
The panel A is the PCA plot of the metabolite profiles from the initial (GLPC/PDF) sample to the Caco-2 transport model. Panel B uses boxplots to show the distribution of the (%) contribution of the compounds belonging to major metabolite groups. Flavs = Flavanoids/coumarins, Cinnamins = Cinnamic acids, Actop = Acetophenones. Boxplots in blue mark represents Dimension 1, while the boxplot marked in black represents Dimension 2.

In Figure 1 panel A, each point represents the metabolite profile measured at each phase of the digestion model. The metabolite profile representing the points for the basolateral fractions of GLPC and PDF very nearly converge, with a distance of 0.3 along the Dimension 1. In contrast the metabolite profile representing the points for the apical fractions of GLPC and PDF are further apart at a distance of 2.4 along Dimension 1. This suggests that irrespective of the disparity in the composition of the metabolites in the initial fractions (separated by a distance of 14.0), the bioavailability of the compounds transported across the Caco-2 monolayer may be limited. This model indicates that much of the metabolites from GLPC/PDF are delivered to the colon for further microbial fermentation.

Metabolite groups such as lignans, flavanoids/coumarins, cinnamic acids, benzoic acids, and acetophenones showed high contribution to the PCA model, particularly along Dimension 1 (Figure 1, panel B). A mass balance of the compounds contributing the highest to the PCA model in Figure 1 is given in Table 1.

**Table 1:** Mass balance of metabolites across initial sample to Caco-2 transportation phase.

| Group | Compound | Sample | Start |  | Salivary | Gastric |  | Intestinal | Apical | Basolateral |
| --- | --- | --- | --- | --- | --- | --- | --- | --- | --- | --- |
| Lignans | secoisolariciresinol | GLPC | 0.18± | 0.08 | 0.15±0.00 | 0.22± | 0.00 | 0.15± 0.00 | 0.03±0.00 | N/D |
|  |  | PDF | 4.10± | 1.68 | 2.88±0.30 | 2.88± | 0.30 | 3.45± 0.08 | 0.92±0.04 | 0.04±0.01 |
|  | lariciresinol | GLPC | 0.21± | 0.37 | 0.36±0.02 | 0.53± | 0.21 | N/D | N/D | N/D |
|  |  | PDF | 7.12± | 2.56 | 4.74±0.37 | 7.92± | 0.48 | 5.67± 0.32 | 0.66±0.57 | N/D |
|  | syringaresinol | GLPC | 17.13± | 8.94 | 15.56±1.23 | 19.21± | 1.95 | 16.58± 0.31 | 1.50±0.06 | N/D |
|  |  | PDF | 402.67± | 179.43 | 321.13±5.93 | 430.54± | 35.71 | 333.67±37.23 | 50.53±2.34 | 18.47±0.54 |
| Indoles | indole-3-carboxylic acid | GLPC | N/D |  | 0.02±0.00 | 0.05± | 0.00 | N/D | N/D | N/D |
|  |  | PDF | 0.30± | 0.13 | 0.21±0.02 | 0.33± | 0.00 | 0.26± 0.00 | 0.08±0.00 | N/D |
| Flavanoids/<br>Coumarins | quercetin-3-glucoside | GLPC | 0.23± | 0.07 | 0.28±0.04 | 0.43± | 0.01 | 0.36± 0.02 | 0.09±0.00 | 0.05±0.00 |
|  |  | PDF | 2.50± | 1.22 | 1.79±0.15 | 2.90± | 0.27 | 1.72± 0.13 | 0.62±0.03 | N/D |
|  | glycitein | GLPC | 1.05± | 0.53 | 1.50±0.01 | 2.25± | 0.07 | 1.32± 0.06 | 0.07±0.01 | 0.03±0.00 |
|  |  | PDF | 12.41± | 7.98 | 11.30±0.30 | 14.16± | 3.17 | 12.18± 0.35 | 0.94±0.08 | 0.69±0.02 |
| Benzoic acids | syringin | GLPC | 2.41± | 1.22 | 2.04±0.13 | 2.95± | 0.14 | 2.25± 0.06 | 0.07±0.06 | N/D |
|  |  | PDF | 22.55± | 6.97 | 16.77±0.41 | 27.20± | 3.07 | 17.76± 1.78 | 0.96±0.22 | 1.56±0.07 |

Values expressed in mean±standard deviation (µg/2g GLPC/PDF).
N/D = not detected.

Data in Table 1 and **Supplementary Table 4**, suggest that the compound groups of indoles, lignans, flavonoids/coumarins, and benzoic acids that were found in high quantities in the PDF fraction, were transported across the Caco-2 monolayer into the basolateral fraction. Individual compounds, namely secoisolariciresinol, lariciresinol, syringaresinol, indole-3-carboxylic acid, quercetin-3-glucoside, glycitein and syringin, showed the most variance and drove the characterisation of the metabolite profile noted in Figure 1.

#### 3.1.2 METABOLITES MEASURED FROM THE CACO-2 APICAL FRACTION TO FAECAL FERMENTATION STAGE

A PCA was performed (Figure 2) using data from **Supplementary Table 3** to study the variance in the metabolites from the apical fraction of the Caco-2 monolayer to the faecal fermentation phase.

**Figure 2:**
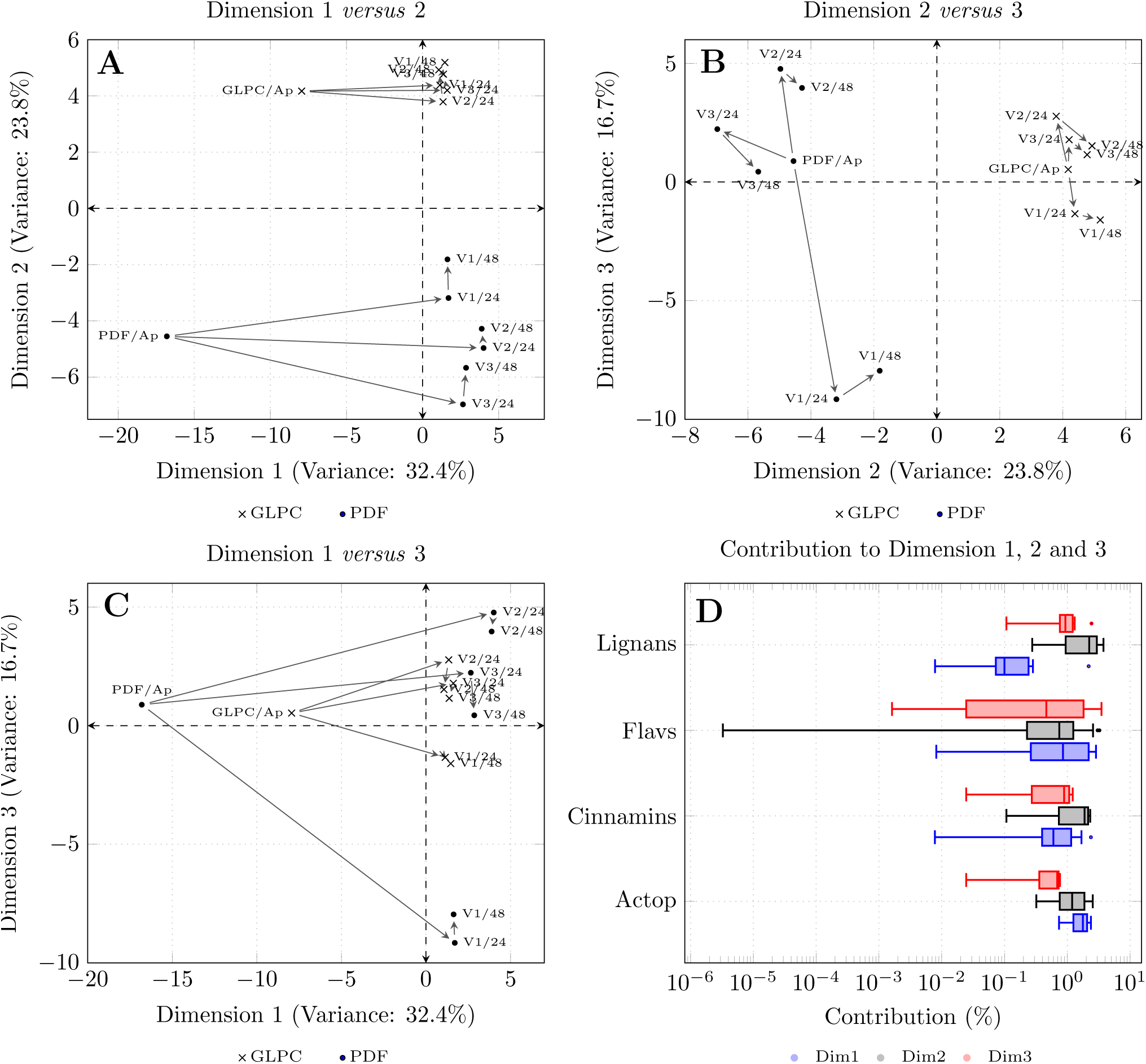
The panels A, B and C show the PCA models from the perspective of Dimensions 1 and 2, Dimensions 2 and 3, and Dimensions 1 and 3 respectively. Panel D shows the distribution of the (%) contribution of the compounds belonging to major metabolite groups. Flavs = Flavanoids/ coumarins, Cinnamins = Cinnamic acids, Actop = Acetophenones, PDF/AP=Protein Depleted Fraction from apical phase of the Caco-2 transport setup, GLPC/AP=Gorse leaf protein concentrate from apical phase of the Caco-2 transport setup. Boxplots in blue mark represents Dimension 1, the boxplot marked in black represents Dimension 2, and the boxplot marked in red represents Dimension 3.

In Figure 2, the panels visualise the PCA model with combinations of Dimensions 1, 2 and 3. Plots showing Dimensions 1 and 2, and Dimensions 1 and 3 place the apical phases of the GLPC/PDF digestates diametrically opposite their respective 24 h and 48 h fermentate metabolite profiles. When considering Dimensions 1 and 2, and Dimensions 2 and 3, there was a clear distinction between metabolites arising from GLPC and PDF for all three volunteer samples. The contribution of individual compounds in each dimension is provided in **Supplementary Code (code chunk 12 and 13)**. The mass balance of highest contributing compounds to the PCA model across Dimensions 1 to 3 is provided in Table 2.

**Table 2:**
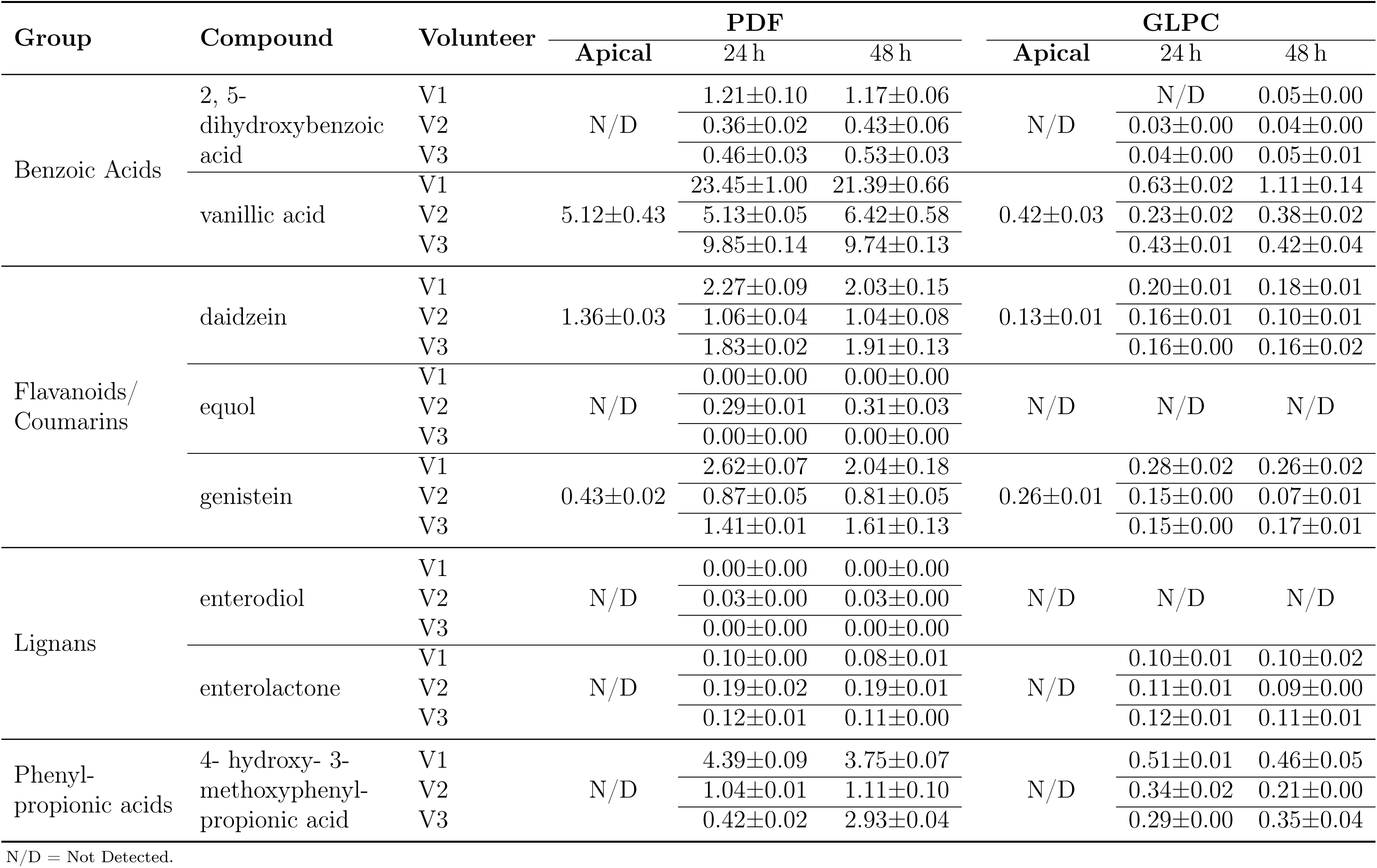
Mass of notable compounds in the faecal fermentation (µg/2g) GLPC/PDF.

| Group | Compound | Volunteer | PDF |  |  | GLPC |  |  |
| --- | --- | --- | --- | --- | --- | --- | --- | --- |
|  |  |  | Apical | 24 h | 48 h | Apical | 24 h | 48 h |
| Benzoic Acids | 2, 5-dihydroxybenzoic acid | V1 | N/D | 1.21±0.10 | 1.17±0.06 | N/D | N/D | 0.05±0.00 |
| | | V2 | | 0.36±0.02 | 0.43±0.06 | | 0.03±0.00 | 0.04±0.00 |
| | | V3 | | 0.46±0.03 | 0.53±0.03 | | 0.04±0.00 | 0.05±0.01 |
| | vanillic acid | V1 | 5.12±0.43 | 23.45±1.00 | 21.39±0.66 | 0.42±0.03 | 0.63±0.02 | 1.11±0.14 |
| | | V2 | | 5.13±0.05 | 6.42±0.58 | | 0.23±0.02 | 0.38±0.02 |
| | | V3 | | 9.85±0.14 | 9.74±0.13 | | 0.43±0.01 | 0.42±0.04 |
| Flavanoids/<br>Coumarins | daidzein | V1 | 1.36±0.03 | 2.27±0.09 | 2.03±0.15 | 0.13±0.01 | 0.20±0.01 | 0.18±0.01 |
| | | V2 | | 1.06±0.04 | 1.04±0.08 | | 0.16±0.01 | 0.10±0.01 |
| | | V3 | | 1.83±0.02 | 1.91±0.13 | | 0.16±0.00 | 0.16±0.02 |
| | equol | V1 | N/D | 0.00±0.00 | 0.00±0.00 | N/D | N/D | N/D |
| | | V2 | | 0.29±0.01 | 0.31±0.03 | | N/D | N/D |
| | | V3 | | 0.00±0.00 | 0.00±0.00 | | N/D | N/D |
| | genistein | V1 | 0.43±0.02 | 2.62±0.07 | 2.04±0.18 | 0.26±0.01 | 0.28±0.02 | 0.26±0.02 |
| | | V2 | | 0.87±0.05 | 0.81±0.05 | | 0.15±0.00 | 0.07±0.01 |
| | | V3 | | 1.41±0.01 | 1.61±0.13 | | 0.15±0.00 | 0.17±0.01 |
| Lignans | enterodiol | V1 | N/D | 0.00±0.00 | 0.00±0.00 | N/D | N/D | N/D |
| | | V2 | | 0.03±0.00 | 0.03±0.00 | | N/D | N/D |
| | | V3 | | 0.00±0.00 | 0.00±0.00 | | N/D | N/D |
| | enterolactone | V1 | N/D | 0.10±0.00 | 0.08±0.01 | N/D | 0.10±0.01 | 0.10±0.02 |
| | | V2 | | 0.19±0.02 | 0.19±0.01 | | 0.11±0.01 | 0.09±0.00 |
| | | V3 | | 0.12±0.01 | 0.11±0.00 | | 0.12±0.01 | 0.11±0.01 |
| Phenyl-propionic acids | 4- hydroxy- 3-methoxyphenyl-propionic acid | V1 | N/D | 4.39±0.09 | 3.75±0.07 | N/D | 0.51±0.01 | 0.46±0.05 |
| | | V2 | | 1.04±0.01 | 1.11±0.10 | | 0.34±0.02 | 0.21±0.00 |
| | | V3 | | 0.42±0.02 | 2.93±0.04 | | 0.29±0.00 | 0.35±0.04 |

N/D = Not Detected.

#### 3.1.3 PCA INCLUDING ALL PHASES OF THE DIGESTION MODEL

The metabolites across the entire *in vitro* digestion model were analysed within a single PCA model to identify compounds driving the categorisation of the metabolite profile in each phase of the digestion. This PCA model is shown in **Supplementary Figure 2** and the contribution of each variable (metabolite) is provided in **Supplementary Code (code chunk 7)**. Major compounds driving categorisation were: histamine, putresine, 4-ethylphenol, 3-methyl-indole, protocatachaldehyde, enterodiol and equol. However, the cos^2^ values of the points representing the metabolite profile in the colonic fermentation phase was <0.5 indicating poor representation in the model.

### 3.2 PERMEABILITY ACROSS THE CACO-2 CELL CULTURE MODEL

#### 3.2.1 TRANSPORT USING STANDARD COMPOUNDS

The permeability of standard compounds across the Caco-2 model was tested individually, as well as an equimolar (50 mM each compound) mixture and the P_app_ values are provided in Table 3. This was performed to check if the permeability of these abundant compounds was additive (i.e. independent of the presence of other compounds) or non-additive (i.e. synergistic, inhibitory, etc.).

**Table 3:** Apparent permeability (*×* 10^−6^ cm^−1^·s^−1^) of standard compounds either as single compounds or a 50 mM equimolar mixture.

| Compound | Single |  |  | Mixed |  |  |
| --- | --- | --- | --- | --- | --- | --- |
| | $P_{\text{app}} (\text{A} \rightarrow \text{B})$ | $P_{\text{app}} (\text{B} \rightarrow \text{A})$ | Ratio | $P_{\text{app}} (\text{A} \rightarrow \text{B})$ | $P_{\text{app}} (\text{B} \rightarrow \text{A})$ | Ratio |
| mandelic acid | 1.28±0.03 | 0.19±0.02 | 23.10 | 7.46 | NA | - |
| chlorogenic acid | 0.44±0.01 | NA | - | 4.33 | NA | - |
| sinapic acid | 1.11±0.01 | 1.75±0.26 | 0.60 | 13.13 | NA | - |
| kaempferol | 4.36±0.26 | NA | - | NA | NA | - |
| p-hydroxybenzoic acid | 1.34±0.08 | 2.42±3.62 | 0.55 | 17.96 | 0.38±0.06 | 47.26 |
| ferulic acid | 8.43±0.03 | 17.61±2.55 | 0.48 | 16.42 | 2.34±0.09 | 7.02 |
| acetovanillone | 15.18±0.59 | 35.58±0.48 | 0.43 | 10.47 | 24.74±0.41 | 0.42 |
| protocatechuic acid | 1.96±0.03 | NA | - | 14.31 | NA | - |
| vanillin | 19.59±0.31 | 38.29±0.16 | 0.51 | NA | NA | - |
| p-coumaric acid | 19.11±0.25 | 15.01±1.01 | 1.27 | 11.91 | 1.07±0.05 | 11.13 |

Values expressed in mean ± standard deviation $\times 10^{-6} \text{ cm}^{-1} \cdot \text{s}^{-1}$ .
Ratio = $\frac{P_{\text{app}} (\text{A} \rightarrow \text{B})}{P_{\text{app}} (\text{B} \rightarrow \text{A})}$ .
N/D=not detected.
Values marked with \* indicate statistically similar $P_{\text{app}}$ values in single compounds as well as mixtures using Student's t-test.

In Table 3, chlorogenic acid, kaemperfol and protocatechuic acid showed non-finite permeability ratios either as a single compound or a mixture. This suggests that efflux proteins governs the active and unidirectional transport of these compounds (Farrell et al., 2012). In the case of acetovanillone, the ratio remained constant for individual and mixed conditions suggesting the action of the efflux proteins on this compound as well. Moreover, it was the only compound with similar P_app (A→B)_ values in single and mixed compound conditions suggesting that its permeability is additive. In the case of p-hydroxybenzoic acid, the P_app (B→A)_ values were statistically similar suggesting that permeability along that direction was also additive. Interestingly though, the P_app (A→B)_ of vanillin and kaempferol was inhibited when compounds were mixed.

#### 3.2.2 TRANSPORT OF GLPC/PDF DIGESTA

The P_app_ values of compounds in the PDF and GLPC experimental digesta are given in **Supplementary Table 6**. Among the standard compounds, only ferulic acid (only in PDF P_app (A→B)_ = 11.89±0.38×10^−6^ cm^−1^·s^−1^) and p-hydroxybenzoic acid (PDF P_app (A→B)_ = 2.34±0.55×10^−6^ cm^−1^·s^−1^; GLPC P_app (A→B)_ = 2.62±1.55×10^−10^ cm^−1^·s^−1^) could be detected in the digesta, and showed transport in at least one direction. In both cases, the P_app_ values were significantly different from the values in Table 3, albeit closer to the single compound setup. This suggests that transport of compounds in the digesta across the Caco-2 layer were synergistic/competitive . Furthermore, a one-way ANOVA on the permeability ratio between the GLPC and PDF indicated significant difference for the same compound, (F_(1,7)_ = 11.39, p = 0.012; excluding single direction transport). This suggests a matrix effect on the permeability of these compounds.

Other compounds of interest identified in the digesta profile were daidzein, genistein, apigenin and vitexin which demonstrated low P_app (A→B)_, but high P_app (B→A)_ values and thus are expected to be confined to the lumen. Similarly, although daidzein and genistein may be associated to beneficial health outcomes (Poschner et al., 2017), they are more likely to remain in the lumen and be delivered to the colon where they are metabolised by the gut microbiota.

### 3.3 TOXICITY OF PLANT METABOLITES TO CACO-2 CELLS

The data provided in the **Supplementary Table 7** shows no significant loss in cell survival for the experimental extracts and their respective digesta. Cell viability did not significantly drop lower than 80% for any treatment, except for camptothecin which was the toxicity-positive control. The tolerance limit of 80% was set by precedence (Farrell et al., 2012) where a significant drop in viability, in tandem with low TEER, suggested to the loss of the monolayer integrity.

The mean cell viability for the pure compounds as well as the digesta was 94.7±13.0% which was not significantly lower than 100% viability using one sample t-test (t = -2.24, df= 29, p= 0.033). For the initial GLPC/PDF samples, the mean viability was 87.1±14.4% and their corresponding digesta showed 84.9±2.8%. A one sample t-test for the Caco-2 viability when exposed to GLPC/PDF digesta was significantly lower than 100% (t = -17.18, df= 9, p= 3.45×10^−8^), but higher than 80% (t= 5.51, df= 9, p= 3.76×10^−4^) as well. Toxicity may have been contributed by the enzymes-only control used in the experiment which showed a Caco-2 viability of 28.2±0.5%. This suggests that PDF and GLPC did not adversely affect cell viability in the Caco-2 model. Analyses are shown in **Supplementary Codes (code chunk 16)**.

### 3.4 SHORT CHAIN FATTY ACIDS (SCFA) PROFILE

The short chain fatty acid content in the faecal fermentation stage was measured at 0 h, 24 h and 48 h (**Supplementary Table 8**). To understand the overall trends in the SCFA profile, a PCA (Figure 3) was performed. The codes and annotations are provided in **Supplementary Codes (code chunk 14)**.

**Figure 3:**
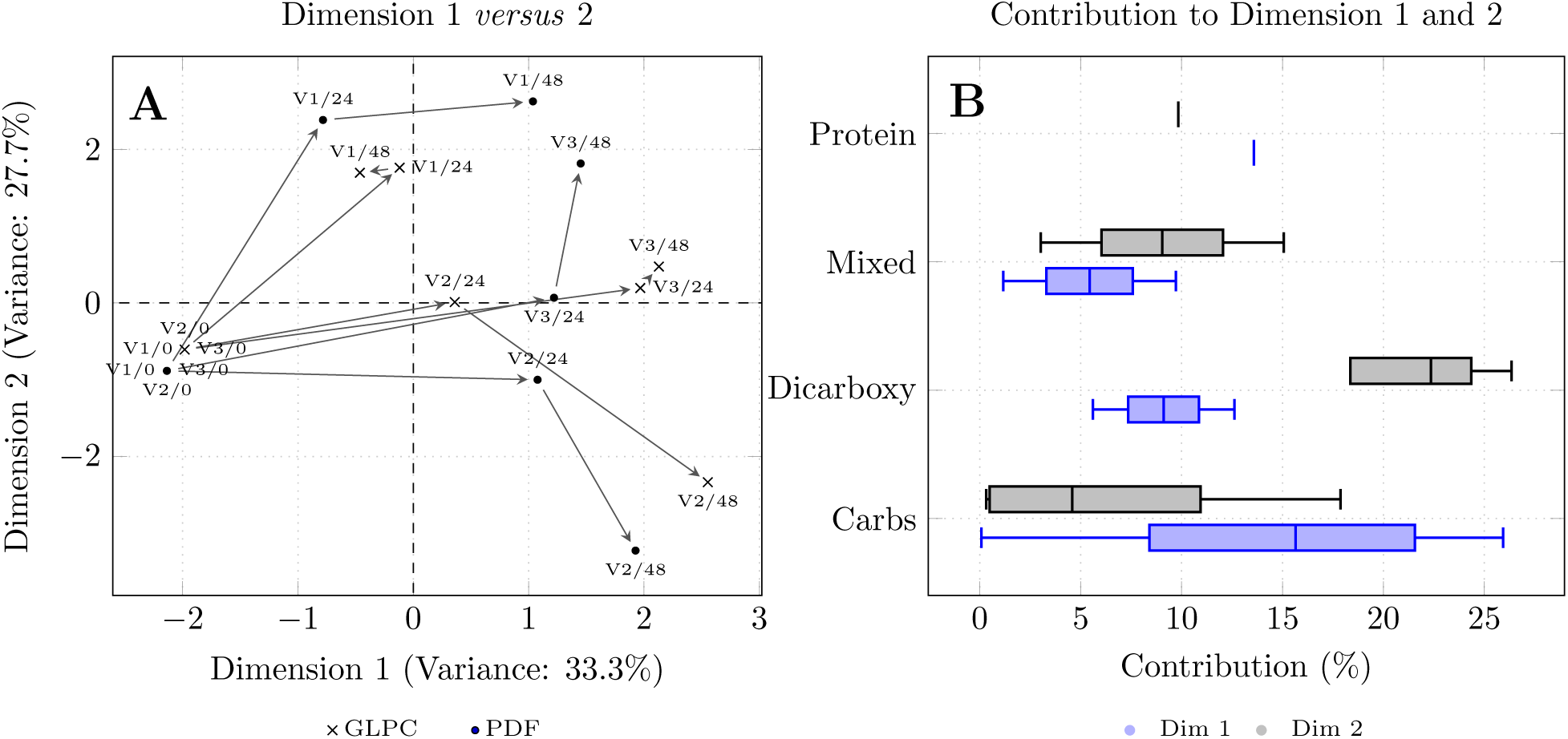
PCA profile of the short chain fatty acids (SCFA) measured in the colonic phase of the in vitro digestion model. The left panel shows the PCA profile of the SCFA at different fermentation times for each volunteer. In panel B, SCFAs are grouped according to the substrates from which they are produced (or dicarboxylic acids that are metabolic intermediates), and their contribution to the PCA profile in the left panel. Carbs=carbohydrates, Dicarboxy=dicarboxylic acids, Mixed= either protein or carbohydrates may be the substrate for the SCFA. Boxplots in blue mark represents Dimension 1, while the boxplot marked in black represents Dimension 2.

Dimension 1 of the PCA model (Figure 3 panel B, blue boxplots) is mainly influenced by the bacterial metabolism of carbohydrates and proteins, while the separation along Dimension 2 was driven by dicarboxylic acids (Figure 3, panel B, black boxplots).

In Figure 3 panel B, the median contribution of SCFAs arising from bacterial metabolism of carbohydrates and protein digestion was high for dimension 1 (marked in blue boxplot), while for dimension 2, dicarboxylic acids showed the highest contribution (marked in black boxplot). In the left panel, the relative placement of the points representing the SCFA profile indicates the changes as colonic fermentation occurs over 48 hours. While the points representing the 48 h profile for all volunteers moved away from their starting point, they differentiate among each other along dimension 2. This suggests that dicarboxylic concentrates drives categorisation between the individual volunteers.

The microbiota of volunteer 2 produced significantly higher butyrate (14.62 mg) compared to volunteer 1 (1.93 mg) and 3 (3.89 mg) at the 48 h time point in the presence of the PDF fraction (**Supplementary Table 8**). Conversely, the microbiota from volunteer 2 showed lower levels of dicarboxylic acids (succinate+lactate = 15.99 mg) compared to volunteer 1 (49.58 mg) and 3 (33.02 mg) after 48 h of colonic fermentation. These trends were similar in the presence of the GLPC fractions as well.

## 4 DISCUSSION

Simulated digestion models provide a controlled and physiologically relevant environment for studying the interactions and evaluating the food safety of novel foods or additives, eliminating the need for animal or human experiments. Leaf protein concentrates from Gorse (GLPC) and its protein depleted fraction (PDF) were subjected to simulated *in vitro* digestion to study the metabolites released and produced as they passed through the alimentary canal.

While previous studies have focused on the digestibility of leaf protein concentrates and used models largely to demonstrate the bio-availability of the protein (Kaur et al., 2021), the work here explored the wider suite of metabolites generated across the digestion process in an attempt to establish benignity. Although recent studies by Prado Massarioli et al. (2023) have used similar *in vitro* digestion models to study the resultant metabolites, the use of anaerobic conditions in this work allowed for closer replication of physiological conditions, and aided *in situ* preservation of metabolites, otherwise lost to oxidation.

### 4.1 ENZYMATIC DIGESTION ENHANCES RELEASE OF METABOLITES FROM THE FOOD MATRIX

Based on the mass balance provided in **Supplementary Table 4**, the recovery of the metabolite groups transported to the basolateral phase by the Caco-2 monolayer (indicating their uptake) shows a relatively high value for benzoic acid (12.9%) with flavonoids/coumarins in a distant second with (7.1%) for GLPC. In the case of PDF, cinnamic acids showed the highest recovery in the intestinal phase at 207.5% relative to the initial content. This may be a result of their release from the matrix of residual proteins. For example, phenyllactic acid was high in the basolateral phase of the PDF digesta (424.1±8.4 µg), showing a 6*×* increase in recovery compared to the initial PDF sample (68.9±0.2 µg) as given in **Supplementary Table 3**. Phenyllactic acids are plant auxins that are phenylalanine catabolites. The Caco-2 cell line used to model the intestinal enterocyte, does not express the necessary enzymes such as phenylalanine hydroxylase (PAH) to produce phenylpyruvic acids (Lichter-Konecki et al., 1999), that is subsequently catabolised to phenyllactic acids. It is likely that freeze-drying and re-dissolution of the PDF intestinal phase digesta in Dulbecco medium for the Caco-2 setup, may have released phenyllactic acids from the matrix.

### 4.2 RESURGENCE OF PLANT METABOLITES IN COLONIC FERMENTATES

The quantities of flavonoids/ coumarins, benzoic acids, and cinnamic acids dropped significantly post exposure to the Caco-2 monolayer (**Supplementary Table 2** and **5**). The combined quantities of compounds in the apical and basolateral fractions were lower than that measured in the preceding intestinal phase. The quantities of these compound groups rose again during the 48 h faecal fermentation (accounting for quantities measured in the controls). This observation may be explained by conjugation of compounds (glucuronidatin, sulphation, methylation) by the Caco-2 monolayer used to simulate the intestinal lining, and the subsequent deconjugation by microbial action during fermentation.

The glucuronidation of xenobiotic compounds by the Caco-2 cell line is well documented and the mechanisms triggering the relevant biochemical systems have been detailed previously by Abid et al. (1995). Likewise, the modification of dietary phenolics by the colonic microbes has been described by Russell et al. (2013), and remains a topic of further investigation.

### 4.3 PERMEABILITY OF THE CACO-2 MONOLAYER WAS SENSITIVE TO COMPOSITE SAMPLES

The rate of transport of compounds across a given surface area (transport velocity), is known as apparent permeability (P_app_). The ratio of P_app(A→B)_ to P_app(B→A)_ indicates the nature of metabolite interaction with the cell surface receptors. A ratio of <1 indicates interaction of metabolites with the efflux transporters (Turner et al., 2000). In other words, the Caco-2 monolayer actively attempts to stop the movement of these metabolites from the apical phase to the basolateral. A ratio of >1 suggests either higher uptake or efflux activity ensuring metabolites do not permeate from the basolateral into the apical phase (Farrell et al., 2012; Čvorović et al., 2018).

The general trend noted in Table 3, was that for a given compound, the P_app (A→B)_ was higher in mixed compound conditions compared to single compounds. Similarly, P_app(B→A)_ dropped below detection levels in mixed conditions compared to single standards. Thus while P_app(A→B)_ value for chlorogenic acid was comparable to that previously observed by Hithamani et al. (2017), its bioavailability from diet is expected to be significantly different.

This observation aligns well with the expected increase in the expression of efflux transporters in response to increased phenolic load (Huang et al., 2008), and that the chosen standard compounds were non-conjugated phytochemicals. The review by Williamson et al. (2018) provides a good description of the fate of dietary phenolics in the human body. Non-conjugated phenolic entities are transported across the intestinal lining, where first conjugations occur, into the blood from where they are transported to the liver and are conjugated further. These conjugated moieties are transported back into the alimentary canal where they are further metabolised by the gut microbiota.

To replicate this effect *in vitro*, the basolateral fraction of the Caco-2 monolayer can be exposed to the HepG2 cell line followed by a re-exposure to the Caco-2 monolayer to study transport of the conjugated moieties from the basolateral fraction to the apical fraction. Unfortunately, this arm of the experiment could not be executed owing to limitations faced during the pandemic. Nonetheless within the performed experiments, partial dynamics of the major compounds identified in Gorse (Iyer et al., 2021) could be examined.

### 4.4 MOST METABOLITES IN THE DIGESTA WERE DELIVERED TO THE COLON

The PCA of the metabolite profile across the entire digestion model is shown in **Supplementary Figure 2**. The cos^2^ values were below even the lenient threshold of 0.5 for the profiles representing the colonic fermentation phase of the model (high quality models generally filter above 0.85). The cos^2^ value is a measure of how well an individual is represented in the PCA model and consequently, how well its variance is captured in the model. By bifurcating the model into non-bacterial and bacterial phases, the metabolite profile of the digesta in each phase is better represented in the model and more internally comparable. The microbes in the colonic fermentation phase have a far more diverse metabolic capability compared to the enzymatic treatment and the mammalian Caco-2 cell line and thus may be prudent to analyse these results independently.

In Figure 1, as GLPC and PDF pass through the digestion model, their overall metabolite profile begins to converge at the basolateral phase of the Caco-2 model. There is low variance between the transported metabolites for PDF and GLPC digesta, suggesting that irrespective of how variable the starting phenolic profile was, the metabolites transported across the Caco-2 monolayer is highly regulated and restrictive. On the other hand the profiles of the apical phase (which represent the fraction expected to stay in the lumen) showed greater variance between each other. Most ingested plant metabolites are thus expected to be delivered to the colon.

### 4.5 BENEFICIAL MICROBIAL METABOLITES WERE NOTED IN FAECAL FERMENTATES

In the faecal fermentation phase of the digestion model, for all volunteers, bacterial metabolism of the phytochemicals such as pinoresinol and secoisolariciresinol to enterolignans such as enterodiol and enterolactone was observed. These metabolites are known to contribute towards beneficial effects, although this model may not truly represent the scale of production and uptake in the human body. Another case is the conversion of daidzein to equol that was noted only in the case for volunteer 2. Despite accounting for the vanillin from the diet, an excess was noted in the supernatants. This may be an indication of the conversion of protocatachaldehyde to vanillin through microbial action.

### 4.6 SCFA PROFILE MAY COMPLIMENT THE PHYTOCHEMICAL METABOLITE PROFILE

The values noted for SCFA were based on pooled fermentate samples. Thus, while statistical comparison was not possible, their PCA profile could provide some indication of the overall trends in the bacterial metabolite trends. Formate, acetate, propionate and butyrate were SCFAs arising from carbohydrate digestion. Valerate was a product of protein metabolism. Iso-butyrate and iso-valerate have mixed substrates and may arise either from carbohydrate and protein metabolism. Lastly, dicarboxylic acids are metabolic intermediates. As shown in panel A of Figure 3, Dimension 1 was largely influenced by carbohydrate and protein metabolites while Dimension 2 was largely influenced by the dicarboxylic acids. Complimenting the phenolic metabolites noted previously, the SCFA profile of the fermentates for volunteer 2 was characterised by low intermediates such as dicarboxylic acids and higher carbohydrate metabolites; particularly butyrate. Correspondingly, the microbiota of volunteer 2 was capable of converting daidzein to equol unlike volunteer 1 and volunteer 3.

### 4.7 DIGESTA SHOWED LOW TOXICITY TO CACO-2 CELLS

Cellular toxicity was measured as a function of the viability of proliferative Caco-2 cells. In the digesta samples noted in **Supplementary Table 7**, there was no case of overt toxicity noted when exposed to Caco-2. Any toxicity measured was largely associated to the enzymes used in the digestion model rather than the extract digesta.

## 5 CONCLUSIONS

The absence of prior reports on toxicity from Gorse, the absence of overt toxicity in proliferative Caco-2 cells, and the absence of any overtly toxic metabolites noted across the simulated digestion model suggests that the gorse leaf protein concentrates (GLPC) and metabolite-rich protein depleted fraction (PDF) may be safe for human consumption.

A majority of the Gorse bioactives are confined to the apical fraction of the digestion model and are expected to be delivered to the colon. While a highly composite bioactive profile appears to enhance permeability for most compounds, it is further modulated by the sample matrix (*viz.* gorse proteins). Gut microbial composition could potentially synergistically enhance health benefits through the production of enterolactones and equol.

## SUPPLEMENTARY MATERIALS

**OSF respository link:** All supplementary materials may be found at the OSF respository at https://osf.io/9qtyz/

**Supplementary Codes**: The codes and output for all statistical analyses performed in R is provided in the file Supplementary Code.pdf.

**Supplementary Materials:** The supplementary materials (Supplementary.pdf file) contains the the Supplementary Figures 1 and 2, and caption for all the Supplementary Tables.

**Supplementary Table 1:** The Supplementary Table 1 (Supplementary_ Table1.csv) shows the chemical names used for in house identification and data processing, and their corresponding trivial names and compound groups.

**Supplementary Table 2:** The Supplementary Table 2 (Supplementary_ Table2.csv) shows the metabolites measured in the fractions (pg·µL^−1^) in the *in vitro* digstion model from the initial GLPC/PDF to the Caco2 monolayer.

**Supplementary Table 3:** The Supplementary Table 3 (Supplementary_ Table3.csv) shows the metabolites measured in the fractions (pg·µL^−1^) in the *in vitro* digstion model from the initial GLPC/PDF to the Caco2 monolayer.

**Supplementary Table 4**: Groupwise summation of compounds (µg/2g) of initial GLPC/PDF to the Caco2 monolayer phase.

**Supplementary Table 5**: Groupwise sum of compounds (µg/2g) of Apical fraction of Caco2 phase to 48 hour colonic fermentation phase.

**Supplementary Table 6**: Apparent permeability of compounds in the Gorse digestates.

**Supplementary Table 7:** The Supplementary Table 7 (Supplementary_ Table7.csv) shows Caco2 viability (%) across experimental fractions at 24 h incubation.

**Supplementary Table 8**: SCFA content of samples across faecal fermentation (mM/5mL).

**Supplementary** **Figure 1**: Process flow of the digestion. Arrows marked in dot-and-dash represent anaerobic setup and those marked in plain represent aerobic setup.

**Supplementary** **Figure 2**: Univariate scaled principal component analysis (PCA) of the metabolite profile measured at each phase of the digestion process of the initial GLPC/PDF.

## Acknowledgements

The authors thank Jenny Martin, Freda Farquharson, and Gillian Donachie for their support with the faecal fermentation experiment. Louise Cantlay helped measure metabolites using the LC/MS. Dr. Teresa Grohmann, Mr. Timo Kramer and Dr. Elaine Hillesheim helped with the correction and editing of the manuscript.

## Funding

This work was funded by the Scottish Government through Rural and Environment Science and Analytical Services Division (RESAS), as part of its strategic funding programme.

## Notes

https://osf.io/9qtyz/

### Competing Interest Statement

The authors have declared no competing interest.

### Summary of Updates

1. Details of the Supplementary text is now shifted to the main manuscript in the Materials and Methods section. 2. Many citations were incorrect. They have now been rectified and updated to APA style citation. 3. Paragraph headers are removed. Numbered sectioning is consistently used.

