## Supplementary_Material for "Simulated digestion, uptake and colonic fermentation of Gorse (*Ulex europeaus*) suggests safe use as a source of protein concentrates and bioactives"

Additional details and codes may be found in the OSF repository: https://osf.io/9qtyz/

#### 1 Supplementary figures

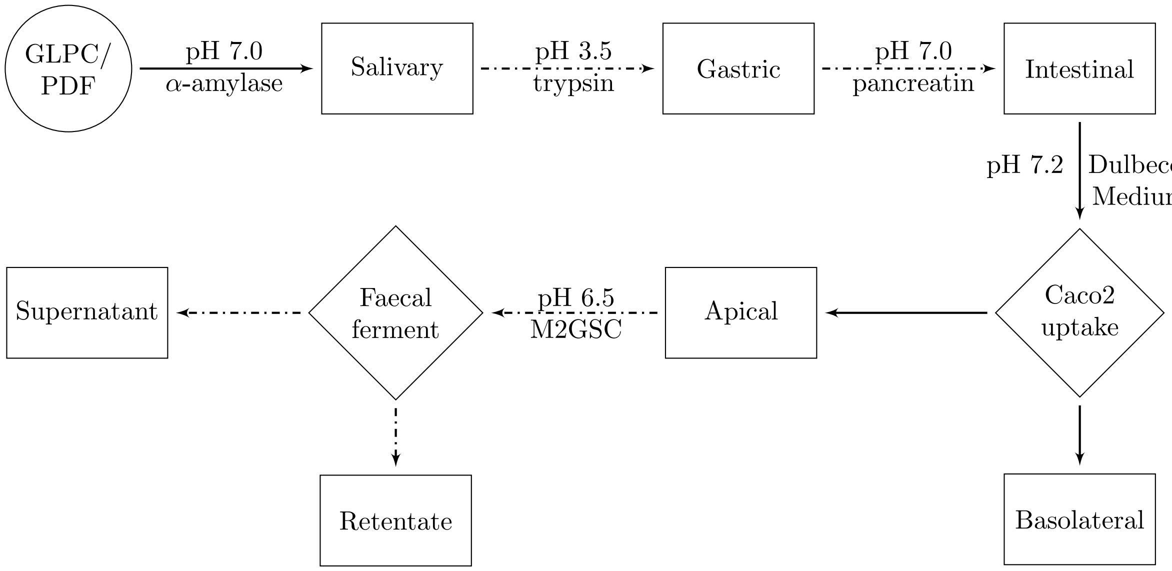


Supplementary table 1: Process flow of the digestion. Arrows marked in dot-and-dash represent anaerobic setup and those marked in plain represent aerobic setup. GLPC = Gorse leaf protein concentrates, PDF = Protein depleted fraction.

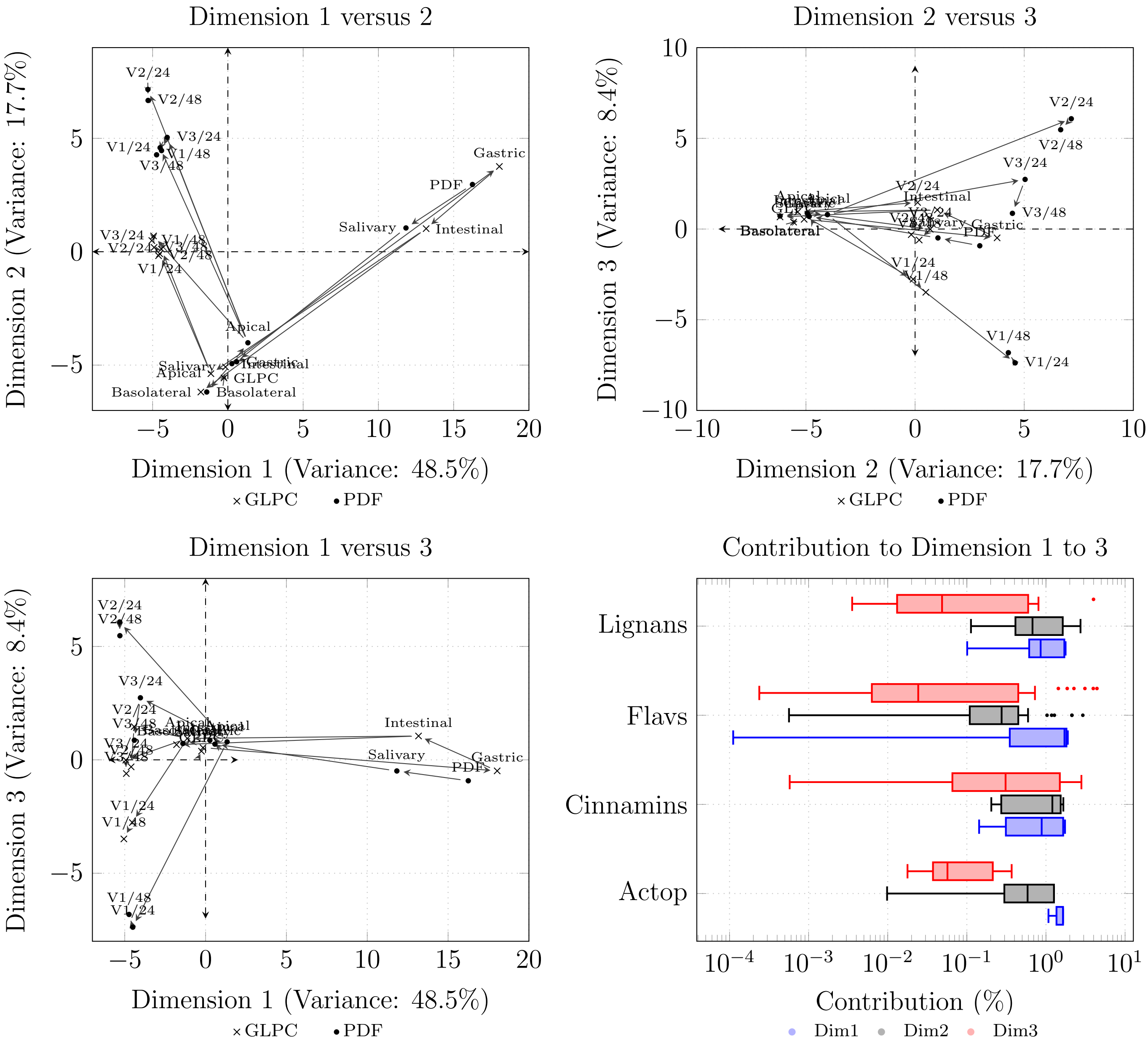


Supplementary table 2: Univariate scaled principal component analysis (PCA) of the metabolite profile measured at each phase of the digstion process of the initial GLPC/PDF.

### 2 Supplementary tables

Supplementary table 1: The chemical names used for in house identification and data processing, and their corresponding trivial names and compound groups.

| This table is available as `Supplementary_Table1.csv` file. |
|---|

[1] Formula is the chemical formula used for in house identification of compounds.
[2] Group refers to the major metabolite group to which the compound belongs.
[3] Trivial is the trivial name given to the compounds for final analysis.

Supplementary table 2: The metabolites measured in the fractions (pg·µL$^{-1}$) in the *in vitro* digstion model from the initial GLPC/PDF to the Caco2 monolayer.

| This table is available as `Supplementary_Table2.csv` file. |
|---|

[1] Sample is either GLPC (Gorse leaf protein concentrate) or PDF (protein depleted fraction). preface by one of the phases, namely, Saliva, Gastric and Intestine indicates the phase in the enzymatic part of the digestion model. Only Saliva, Gastric or Intestine are the controls.
[2] Volunteer is common which is irrelevant in this part of the model. This column is used to distinguish relevant features for data processing when subsequently combining it with the colonic phase of the model.
[3] Part refers to either the enzyme based "Digestion" or the Caco2 model "Transport".
[4] Time is either as "Digestion", the "Supernatant" which refers to the apical fraction of the Caco2 model or numbers from 1 to 4 which are the time points at which the basolateral fractions were collected.
[5] Trial is the replicate number for a given condition.

Supplementary table 3: The metabolites measured in the fractions (pg·µL$^{-1}$) in the *in vitro* digstion model from the initial GLPC/PDF to the Caco2 monolayer.

| This table is available as `Supplementary_Table3.csv` file. |
|---|

[1] Sample is either GLPC (Gorse leaf protein concentrate), PDF (protein depleted fraction) fermentates or their controls, namely, the unfermented GLPC/PDF, the M2GSC media, and the faecal samples of each volunteer.
[2] In Volunteer, Common refers to the common controls in the fermentation stage. It also refers to volunteers 1, 2, and 3 who donated faecal samples for this experiment.
[3] Part refers to "Colonic" as this is the colonic phase of the model.
[4] Time is the fermentation time measured in hours. It is either 0, 24 or 48.
[5] Trial is the replicate number for a given condition.

Supplementary table 4: Groupwise summation of compounds (µg/2g) of initial GLPC/PDF to the Caco2 monolayer phase

| **Group** | **Sample** | **Start** | **Saliva** | **Gastric** | **Intestinal** | **Apical** | **Basolateral** | **%R$_{(Int)}$** | **%R$_{(bas)}$** |
|---|---|---|---|---|---|---|---|---|---|
| Acetophenones | GLPC | 0.12 ± 0.02 | 0.11 ± 0.00 | 0.15 ± 0.00 | 0.07 ± 0.00 | N/D | N/D | 58.33 ± 0.00 | N/D |
| | PDF | 1.75 ± 0.28 | 1.25 ± 0.04 | 2.15 ± 0.05 | 1.35 ± 0.02 | 0.03 ± 0.00 | 0.02 ± 0.00 | 77.14 ± 7.14 | 1.14 ± 0.00 |
| Amines | GLPC | 1.68 ± 0.27 | 5.52 ± 0.07 | 10.19 ± 0.02 | 39.32 ± 1.59 | 40.92 ± 0.80 | 89.85 ± 0.70 | 2340.48 ± 588.89 | 5348.21 ± 2.59 |
| | PDF | 17.11 ± 1.92 | 17.93 ± 0.47 | 46.74 ± 0.49 | 84.24 ± 0.60 | 42.2 ± 9.8 | 94.25 ± 1.06 | 492.34 ± 31.25 | 550.85 ± 0.55 |
| Benzaldehydes | GLPC | 0.07 ± 0.03 | 0.14 ± 0.01 | 0.24 ± 0.01 | 0.18 ± 0.01 | N/D | 0.05 ± 0.01 | 257.14 ± 33.33 | 71.43 ± 0.33 |
| | PDF | 1.92 ± 0.57 | 1.6 ± 0.7 | 2.32 ± 0.15 | 1.52 ± 0.04 | N/D | 0.13 ± 0.01 | 79.17 ± 7.02 | 6.77 ± 0.02 |
| Benzoic acids | GLPC | 6.35 ± 0.56 | 5.34 ± 0.06 | 7.32 ± 0.06 | 6.75 ± 0.08 | 2.78 ± 0.07 | 0.82 ± 0.02 | 106.3 ± 142.9 | 12.91 ± 0.04 |
| | PDF | 101.68 ± 6.73 | 77.48 ± 0.32 | 115.8 ± 12.1 | 82.34 ± 0.87 | 26.5 ± 4.8 | 2.76 ± 0.03 | 80.98 ± 12.93 | 2.71 ± 0.00 |
| Cinnamic acids | GLPC | 1.59 ± 0.15 | 1.35 ± 0.02 | 3.05 ± 0.03 | 3.99 ± 0.02 | 1.82 ± 0.06 | N/D | 250.94 ± 13.33 | N/D |
| | PDF | 23.81 ± 2.87 | 18.32 ± 0.26 | 28.75 ± 0.55 | 49.41 ± 2.19 | 19.10 ± 0.70 | 2.15 ± 0.04 | 207.52 ± 76.31 | 9.03 ± 0.01 |
| Flavanoids/coumarins | GLPC | 14.86 ± 0.61 | 13.56 ± 0.08 | 18.12 ± 0.20 | 14.70 ± 0.08 | 1.81 ± 0.01 | 1.21 ± 0.00 | 98.92 ± 13.11 | 8.14 ± 0.00 |
| | PDF | 105.60 ± 3.66 | 73.24 ± 0.17 | 107.71 ± 0.75 | 72.08 ± 0.20 | 9.38 ± 0.03 | 6.37 ± 0.02 | 68.26 ± 5.46 | 6.03 ± 0.01 |
| Indoles | GLPC | N/D | 0.24 ± 0.00 | 0.45 ± 0.00 | 0.29 ± 0.01 | 0.03 ± 0.00 | 0.03 ± 0.00 | N/A | N/A |
| | PDF | 0.72 ± 0.14 | 0.64 ± 0.01 | 1.05 ± 0.01 | 0.99 ± 0.01 | 0.12 ± 0.00 | 0.04 ± 0.00 | 137.50 ± 7.14 | 5.56 ± 0.00 |
| Lignans | GLPC | 18.15 ± 4.01 | 16.59 ± 0.55 | 20.72 ± 0.87 | 17.44 ± 0.14 | 1.52 ± 0.03 | N/D | 96.09 ± 3.49 | N/D |
| | PDF | 427.17 ± 80.29 | 336.46 ± 2.69 | 457.36 ± 15.97 | 354.40 ± 16.65 | 53.09 ± 1.08 | 19.43 ± 0.24 | 82.96 ± 20.74 | 4.55 ± 0.00 |
| Mandelic acid | GLPC | 1.19 ± 0.07 | N/D | N/D | N/D | N/D | N/D | N/D | N/D |
| | PDF | 2.49 ± 1.38 | 2.11 ± 0.28 | 2.90 ± 0.05 | N/D | N/D | N/D | N/D | N/D |
| Phenolic alcohol | GLPC | 0.85 ± 0.58 | 0.1 ± 0.0 | N/D | N/D | N/D | N/D | N/D | N/D |
| | PDF | 0.73 ± 0.04 | 0.08 ± 0.01 | N/D | N/D | N/D | N/D | N/D | N/D |
| Other phenolics | GLPC | 0.16 ± 0.08 | 0.12 ± 0.01 | 0.24 ± 0.03 | 0.15 ± 0.00 | N/D | N/D | 93.75 ± 0.00 | N/D |
| | PDF | 5.47 ± 3.28 | 4.86 ± 0.24 | 5.69 ± 0.33 | 2.59 ± 0.23 | N/D | N/D | 47.35 ± 7.01 | N/D |
| Phenylacetic acids | GLPC | N/D | 0.08 ± 0.00 | 0.13 ± 0.01 | 0.10 ± 0.00 | 1.05 ± 0.06 | 2.48 ± 0.04 | N/A | N/A |
| | PDF | 6.08 ± 2.19 | 6.77 ± 0.24 | 5.42 ± 0.36 | 1.25 ± 0.07 | 2.58 ± 0.25 | 2.44 ± 0.03 | 20.56 ± 3.20 | 40.13 ± 0.01 |
| Phenyllactic acids | GLPC | N/D | N/D | N/D | N/D | 0.15 ± 0.01 | 0.45 ± 0.01 | N/A | N/A |
| | PDF | 0.07 ± 0.04 | 0.05 ± 0.00 | 0.07 ± 0.00 | 0.07 ± 0.01 | 0.23 ± 0.04 | 0.42 ± 0.01 | 100.00 ± 25.00 | 600.00 ± 0.25 |
| Phenylpropionic acids | GLPC | N/D | N/D | N/D | N/D | N/D | N/D | N/A | N/A |
| | PDF | 1.71 ± 0.62 | 1.37 ± 0.03 | 1.97 ± 0.12 | 1.42 ± 0.09 | N/D | N/D | 83.04 ± 14.52 | N/D |
| Phenypyruvic acids | GLPC | 0.30 ± 0.09 | 0.31 ± 0.02 | 0.67 ± 0.04 | 0.96 ± 0.05 | 1.73 ± 0.31 | 2.22 ± 0.02 | 320.00 ± 55.56 | 740.00 ± 0.22 |
| | PDF | 0.32 ± 0.11 | 0.38 ± 0.01 | 0.93 ± 0.02 | 1.48 ± 0.11 | 1.10 ± 0.05 | 2.86 ± 0.32 | 462.50 ± 100.00 | 893.75 ± 2.91 |

[1] %R$_{(Int)}$ is the recovery of compounds in the intestinal phase relative to the intial GLPC/PDF sample.

[2] %R$_{(bas)}$ is the recovery of the compounds in the basolateral phase of the Caco2 monolayer. In Volunteer, Common refers to the common controls in the fermentation stage. It also refers to volunteers 1, 2, and 3 who donated faecal samples for this experiment.

[3] Part refers to "Colonic" as this is the colonic phase of the model.

[4] Time is the fermentation time measured in hours. It is either 0, 24 or 48.

[5] Trial is the replicate number for a given condition.

Supplementary table 5: Groupwise sum of compounds (µg/2g) of Apical fraction of Caco2 phase to 48 hour colonic fermentation phase

| **Group** | **Sample** | **Apical** | **24 h** | **%R$_{(24)}$** | **48 h** | **%R$_{(48)}$** |
|---|---|---|---|---|---|---|
| Acetophenones | GLPC | N/D | $0.22 \pm 0.01$ | – | $0.31 \pm 0.01$ | – |
| | PDF | $0.03 \pm 0.00$ | $1.71 \pm 0.01$ | 5700.0 | $1.47 \pm 0.01$ | 4900.0 |
| Amines | GLPC | $40.92 \pm 0.80$ | $997.39 \pm 2.39$ | 2437.4 | $1449.72 \pm 6.87$ | 3542.8 |
| | PDF | $42.20 \pm 0.98$ | $783.26 \pm 0.76$ | 1856.1 | $871.05 \pm 1.69$ | 2064.1 |
| Benzaldehydes | GLPC | N/D | $0.35 \pm 0.00$ | – | $0.09 \pm 0.00$ | – |
| | PDF | N/D | $0.48 \pm 0.01$ | – | $0.45 \pm 0.01$ | – |
| Benzenes | GLPC | N/D | N/D | – | N/D | – |
| | PDF | N/D | $6.80 \pm 0.41$ | – | N/D | – |
| Benzoic acids | GLPC | $2.78 \pm 0.07$ | $10.77 \pm 0.07$ | 387.4 | $10.76 \pm 0.05$ | 387.1 |
| | PDF | $26.50 \pm 0.48$ | $77.18 \pm 0.20$ | 291.2 | $75.5 \pm 1.7$ | 284.9 |
| Bile acids | GLPC | N/D | $7.88 \pm 0.16$ | – | $7.40 \pm 0.24$ | – |
| | PDF | N/D | $5.46 \pm 0.07$ | – | $3.53 \pm 0.06$ | – |
| Cinnamic acids | GLPC | $1.82 \pm 0.06$ | $1.98 \pm 0.01$ | 108.8 | $0.05 \pm 0.00$ | 2.7 |
| | PDF | $19.10 \pm 0.70$ | $85.99 \pm 0.38$ | 450.2 | $24.83 \pm 0.30$ | 130.0 |
| Flavanoids/coumarins | GLPC | $1.81 \pm 0.01$ | $2.43 \pm 0.00$ | 134.3 | $2.22 \pm 0.01$ | 122.7 |
| | PDF | $9.38 \pm 0.03$ | $23.14 \pm 0.02$ | 246.7 | $23.09 \pm 0.06$ | 246.2 |
| Indoles | GLPC | $0.03 \pm 0.00$ | $3.99 \pm 0.02$ | 13,300.0 | $4.89 \pm 0.04$ | 16,300.0 |
| | PDF | $0.12 \pm 0.00$ | $4.84 \pm 0.01$ | 4033.3 | $5.79 \pm 0.02$ | 4825.0 |
| Lignans | GLPC | $1.52 \pm 0.03$ | $4.55 \pm 0.07$ | 299.3 | $3.32 \pm 0.06$ | 218.4 |
| | PDF | $53.09 \pm 1.08$ | $108.75 \pm 0.50$ | 204.8 | $97.54 \pm 0.28$ | 183.7 |
| Phenolic alcohol | GLPC | N/D | N/D | – | N/D | – |
| | PDF | N/D | $0.06 \pm 0.00$ | – | $0.06 \pm 0.00$ | – |
| Phenolic dimers | GLPC | N/D | $0.19 \pm 0.00$ | – | $0.17 \pm 0.01$ | – |
| | PDF | N/D | $0.42 \pm 0.02$ | – | $0.39 \pm 0.01$ | – |
| Other phenolics | GLPC | N/D | $17.90 \pm 0.41$ | – | $19.20 \pm 0.85$ | – |
| | PDF | N/D | $12.48 \pm 0.30$ | – | $15.32 \pm 1.23$ | – |
| Phenylacetic acids | GLPC | $1.05 \pm 0.06$ | $33.35 \pm 0.20$ | 3176.2 | $33.76 \pm 0.52$ | 3215.2 |
| | PDF | $2.58 \pm 0.25$ | $35.53 \pm 0.25$ | 1377.1 | $33.39 \pm 0.31$ | 1294.2 |
| Phenyllactic acids | GLPC | $0.15 \pm 0.01$ | $19.92 \pm 0.27$ | 13,280.0 | $36.6 \pm 6.3$ | 24,400.0 |
| | PDF | $0.23 \pm 0.04$ | $59.45 \pm 0.56$ | 25,847.8 | $76.77 \pm 1.01$ | 33,378.3 |
| Phenylpropionic acids | GLPC | N/D | $86.06 \pm 0.39$ | – | $93.24 \pm 1.11$ | – |
| | PDF | N/D | $105.82 \pm 0.48$ | – | $116.68 \pm 0.92$ | – |
| Phenypyruvic acids | GLPC | $1.73 \pm 0.31$ | $2.20 \pm 0.05$ | 127.2 | $0.59 \pm 0.01$ | 34.1 |
| | PDF | $1.10 \pm 0.05$ | $2.68 \pm 0.02$ | 243.6 | $1.36 \pm 0.03$ | 123.6 |

[1] Measurements across volunteers were first averaged, and then summed according to metabolite group.
[2] %R$_{(24)}$ refers to the recovery of the metabolite group after 24 hours of fermentation relative to the apical fraction of the Caco2 monolayer phase.
[3] %R$_{(48)}$ refers to the recovery of the metabolite group after 24 hours of fermentation relative to the apical fraction of the Caco2 monolayer phase.

Supplementary table 6: Apparent permeability of compounds in the Gorse digestates.

| **Compounds** | **PDF** | | | **GLPC** | | |
|---|---|---|---|---|---|---|
| | $\mathbf{P_{app\ (A\rightarrow B)}}$ | $\mathbf{P_{app\ (B\rightarrow A)}}$ | **Ratio** | $\mathbf{P_{app\ (A\rightarrow B)}}$ | $\mathbf{P_{app\ (B\rightarrow A)}}$ | **Ratio** |
| 3,4,5–trimethoxyacetophenone | 6.85±1.35 | B.D.L | N/A | B.D.L | $(4.22 \pm 1.34) \times 10^{-8}$ | 0.00 |
| 4–hydroxyphenylpyruvic acid | 253.20±33.23 | 47.24±2.60 | 5.36 | $(4.83 \pm 0.63) \times 10^{-8}$ | $(9.17 \pm 1.80) \times 10^{-9}$ | 5.26 |
| 5–hydroxytryptophan | 592.53±114.42 | 42.68±9.46 | 13.89 | $(1.01 \pm 0.18) \times 10^{-7}$ | $(2.27 \pm 0.04) \times 10^{-8}$ | 4.43 |
| apigenin | 13.47±6.35 | 28.38±9.82 | 0.48 | B.D.L | $(1.78 \pm 0.22) \times 10^{-8}$ | 0.00 |
| benzoic acid | B.D.L | 45.16±9.64 | 0.00 | B.D.L | $(9.01 \pm 1.88) \times 10^{-9}$ | 0.00 |
| biochanin A | B.D.L | B.D.L | N/A | B.D.L | $(3.56 \pm 0.02) \times 10^{-8}$ | 0.00 |
| daidzein | 12.00±1.51 | 18.08±12.81 | 0.66 | B.D.L | $(6.30 \pm 0.81) \times 10^{-8}$ | 0.00 |
| eriocitrin | B.D.L | 5.61±7.94 | N/A | B.D.L | B.D.L | N/A |
| ferulic acid | 11.89±0.38 | B.D.L | N/A | B.D.L | B.D.L | N/A |
| formononetin | 14.36±1.69 | 29.38±5.37 | 0.49 | $(1.33 \pm 0.18) \times 10^{-9}$ | $(2.33 \pm 0.35) \times 10^{-8}$ | 0.06 |
| genistein | B.D.L | 28.08±0.18 | 0.00 | B.D.L | $(2.31 \pm 0.27) \times 10^{-8}$ | 0.00 |
| glycitein | 6.34±0.73 | 24.85±5.34 | 0.26 | $(4.64 \pm 0.72) \times 10^{-11}$ | $(2.94 \pm 0.50) \times 10^{-8}$ | $1.57 \times 10^{-3}$ |
| indole | B.D.L | 61.89±19.11 | 0.00 | B.D.L | $(2.26 \pm 0.32) \times 10^{-8}$ | 0.00 |
| indole–3–acetonitrile | 12.55±1.71 | B.D.L | N/A | $(1.58 \pm 0.02) \times 10^{-8}$ | B.D.L | N/A |
| indole–3–acrylic acid | 211.38±16.84 | B.D.L | N/A | $(4.23 \pm 0.39) \times 10^{-8}$ | B.D.L | N/A |
| indole–3–carbinol | B.D.L | 46.10±5.31 | 0.00 | B.D.L | $(1.01 \pm 0.08) \times 10^{-8}$ | 0.00 |
| indole–3–carboxaldehyde | B.D.L | 44.82±5.18 | 0.00 | B.D.L | $(8.82 \pm 0.96) \times 10^{-8}$ | 0.00 |
| indole–3–lactic acid | 829.28±108.20 | B.D.L | N/A | $(1.62 \pm 0.17) \times 10^{-7}$ | B.D.L | N/A |
| indole–3–pyruvic acid | B.D.L | 42.86±10.82 | 0.00 | B.D.L | $(9.27 \pm 1.28) \times 10^{-9}$ | 0.00 |
| *iso*–liquiritigenin | 20.64±2.40 | 31.95±3.24 | 0.64 | B.D.L | $(3.02 \pm 0.37) \times 10^{-8}$ | 0.00 |
| kynurenic acid | 8.40±1.36 | 38.80±5.99 | 0.22 | $(1.71 \pm 0.22) \times 10^{-9}$ | $(1.15 \pm 0.15) \times 10^{-8}$ | 0.15 |
| naringenin | 23.97±6.69 | B.D.L | N/A | B.D.L | B.D.L | N/A |
| *p*–hydroxybenzaldehyde | 6.91±1.34 | B.D.L | N/A | $(4.57 \pm 3.72) \times 10^{-10}$ | B.D.L | N/A |
| *p*–hydroxybenzoic acid | 2.34±0.55 | B.D.L | N/A | $(2.62 \pm 1.55) \times 10^{-10}$ | B.D.L | N/A |
| phenylacetic acid | 92.51±8.85 | B.D.L | N/A | $(1.86 \pm 0.34) \times 10^{-8}$ | B.D.L | N/A |
| phenyllactic acid | B.D.L | 45.90±8.61 | 0.00 | B.D.L | $(9.03 \pm 1.83) \times 10^{-9}$ | 0.00 |
| phenylpyruvic acid | B.D.L | 38.20±5.56 | 0.00 | B.D.L | $(1.01 \pm 0.01) \times 10^{-8}$ | 0.00 |
| pinoresinol | 6.26±0.94 | B.D.L | N/A | B.D.L | $(7.81 \pm 1.41) \times 10^{-8}$ | 0 |

Continued on next page

**Table 6 – continued from previous page**

| Compounds | PDF | | | GLPC | | |
|---|---|---|---|---|---|---|
| | $P_{app\ (A\rightarrow B)}$ | $P_{app\ (B\rightarrow A)}$ | Ratio | $P_{app\ (A\rightarrow B)}$ | $P_{app\ (B\rightarrow A)}$ | Ratio |
| piperidine | 305.67±53.47 | 46.03±6.40 | 6.65 | $(5.93 \pm 0.85) \times 10^{-8}$ | $(9.06 \pm 1.20) \times 10^{-9}$ | 6.55 |
| putersine | B.D.L | 36.07±5.15 | 0.00 | B.D.L | $(7.93 \pm 0.94) \times 10^{-9}$ | 0.00 |
| quercetin–3–glucoside | B.D.L | B.D.L | N/A | $(2.10 \pm 1.97) \times 10^{-10}$ | B.D.L | N/A |
| salicylic acid | 11.43±1.31 | B.D.L | 0.00 | $(1.05 \pm 0.74) \times 10^{-9}$ | B.D.L | 0.00 |
| seco–*iso*–lariciresinol | 0.14±0.07 | B.D.L | N/A | B.D.L | $(2.14 \pm 0.29) \times 10^{-8}$ | 0.00 |
| spermidine | 0.79±0.77 | B.D.L | N/A | $(3.40 \pm 0.48) \times 10^{-11}$ | B.D.L | N/A |
| spermine | B.D.L | 53.14±5.55 | 0.00 | B.D.L | $(1.38 \pm 0.02) \times 10^{-8}$ | 0.00 |
| syringaresinol | 3.49±0.58 | B.D.L | N/A | B.D.L | $(4.48 \pm 0.70) \times 10^{-8}$ | 0.00 |
| syringin | 4.79±1.25 | B.D.L | N/A | B.D.L | B.D.L | N/A |
| tangeretin | B.D.L | 62.14±23.71 | 0.00 | B.D.L | $(9.19 \pm 0.64) \times 10^{-9}$ | 0.00 |
| vitexin | B.D.L | 22.14±2.58 | 0.00 | B.D.L | $(2.55 \pm 0.20) \times 10^{-8}$ | 0.00 |

Values expressed in $10^{-6}\ cm^{-1} \cdot s^{-1}$.

B.D.L= $P_{app}$ values equal 0±0 as no compound was detected on across the Caco2 monolayer.

A ratio of 0.00 or N/A= Unidirectional transport or invalid ratio.

Ratio= $\frac{P_{app(A-B)}}{P_{app(B-A)}}$

Supplementary table 7: Caco2 viability (%) across experimental fractions at 24 h incubation.

| |
|---|
| This table is available as `Supplementary_Table7.csv` file. |

[1] Conditions marked 'intestinal' represent the digesta at the intestinal phase of the digestion model.
[2] 'Enzyme only' represents control which contains neither PDF not GLPC. Mandelic acid to p–hydroxybenzoic acid represent standard compounds used at 50 mM concentration.

Supplementary table 8: SCFA content of samples across fæcal fermentation (mM/5mL).

| Volunteer | Sample | Time | Formate | Acetate | Propionate | iso-Butyrate | Butyrate | iso-Valerate | Valerate | Lactate | Succinate |
|---|---|---|---|---|---|---|---|---|---|---|---|
| **Control** | **Only Media** | 0 | N/D | 18.2 | 4.21 | 0.16 | 4.69 | N/D | 0.37 | N/D | N/D |
| | | 24 | N/D | 18.2 | 4.21 | 0.16 | 4.69 | N/D | 0.37 | N/D | N/D |
| | | 48 | N/D | 17.47 | 4.19 | 0.15 | 4.49 | N/D | 0.36 | N/D | N/D |
| | **Only PDF** | 0 | N/D | 18.28 | 4.94 | 0.16 | 4.39 | N/D | 0.35 | N/D | N/D |
| | | 24 | N/D | 17.28 | 3.94 | N/D | 4.39 | N/D | 0.35 | N/D | N/D |
| | | 48 | 0.35 | 17.77 | 4.01 | 0.15 | 4.48 | N/D | 0.35 | N/D | N/D |
| | **Only GLPC** | 0 | N/D | 18.18 | 4.25 | 0.16 | 4.57 | N/D | 0.36 | N/D | N/D |
| | | 24 | 0.41 | 18.18 | 4.25 | 0.16 | 4.57 | 0.11 | 0.36 | 0.46 | N/D |
| | | 48 | N/D | 18.04 | 4.34 | 0.16 | 4.51 | N/D | 0.36 | N/D | N/D |
| **Volunteer 1** | **Only Faeces** | 0 | 0.83 | 18.58 | 4.33 | N/D | 4.68 | N/D | 0.37 | N/D | N/D |
| | | 24 | 8.17 | 28.2 | 4.57 | 0.16 | 4.59 | N/D | 0.37 | 25.21 | 3.86 |
| | | 48 | 5.82 | 31.8 | 6.85 | 0.17 | 4.57 | N/D | 0.36 | 25.35 | 2.56 |
| | **PDF** | 0 | N/D | 18.28 | 4.94 | 0.16 | 4.39 | N/D | 0.35 | N/D | N/D |
| | | 24 | 5.75 | 28.45 | 3.93 | 0.14 | 4.05 | N/D | 0.32 | 45.87 | 7.68 |
| | | 48 | 0.71 | 39.96 | 5.11 | N/D | 4.99 | N/D | 0.38 | 96.34 | 10.48 |
| | **GLPC** | 0 | N/D | 18.18 | 4.25 | 0.16 | 4.57 | N/D | 0.36 | N/D | N/D |
| | | 24 | 5.61 | 28.72 | 3.95 | 0.15 | 4.39 | N/D | 0.35 | 70.25 | 5.43 |
| | | 48 | 2.49 | 31.45 | 4.52 | 0.15 | 4.39 | N/D | 0.34 | 86.54 | 5.37 |
| **Volunteer 2** | **Only Faeces** | 0 | 0.8 | 18.05 | 4.37 | 0.15 | 4.62 | 0.1 | 0.37 | N/D | N/D |
| | | 24 | 9.49 | 22.51 | 7.01 | 0.17 | 32.1 | N/D | 0.43 | 6.7 | 1.18 |
| | | 48 | 7.93 | 29.27 | 9.67 | 0.2 | 40.93 | 0.14 | 0.54 | 6.36 | N/D |
| | **PDF** | 0 | N/D | 18.28 | 4.94 | 0.16 | 4.39 | N/D | 0.35 | N/D | N/D |
| | | 24 | 2.82 | 93.93 | 3.99 | 0.18 | 12.27 | 0.13 | 0.37 | 30.82 | 1.39 |
| | | 48 | 3.36 | 84.03 | 5.31 | 0.18 | 33.18 | N/D | 0.41 | 34.81 | 0.53 |
| | **GLPC** | 0 | N/D | 18.18 | 4.25 | 0.16 | 4.57 | N/D | 0.36 | N/D | N/D |
| | | 24 | 6.91 | 95.65 | 4.08 | 0.17 | 5.12 | N/D | 0.36 | 32.41 | 0.85 |
| | | 48 | 8.87 | 86.85 | 5.12 | 0.16 | 25.55 | 0.12 | 0.38 | 34.27 | 0.73 |
| **Volunteer 3** | **Only Faeces** | 0 | 0.8 | 17.92 | 4.37 | 0.16 | 4.72 | 0.1 | 0.37 | N/D | N/D |
| | | 24 | – | – | – | – | – | – | – | – | – |
| | | 48 | 7.86 | 44.42 | 9.75 | 0.19 | 17.17 | 0.16 | 0.39 | 12.48 | 1.3 |
| | **PDF** | 0 | N/D | 18.28 | 4.94 | 0.16 | 4.39 | 0 | 0.35 | N/D | N/D |
| | | 24 | 4.41 | 105 | 4.28 | 0.16 | 5.59 | 0.11 | 0.36 | 47.58 | 3.17 |
| | | 48 | 4.69 | 111.23 | 4.22 | 0 | 5.69 | N/D | 0.37 | 68.77 | 3.47 |
| | **GLPC** | 0 | N/D | 18.18 | 4.25 | 0.16 | 4.57 | N/D | 0.36 | N/D | N/D |
| | | 24 | 7.61 | 94.19 | 4.25 | 0.16 | 7.65 | 0.12 | 0.37 | 45.42 | 5.41 |
| | | 48 | 10.64 | 107.59 | 4.3 | 0.17 | 8.83 | N/D | 0.38 | 49.68 | 6.05 |

N/D = Not detected.
The 24 h time point for volunteer 3 (only faeces) was lost due to evaporation.
